# ASTRAL-Pro: quartet-based species tree inference despite paralogy

**DOI:** 10.1101/2019.12.12.874727

**Authors:** Chao Zhang, Celine Scornavacca, Erin K. Molloy, Siavash Mirarab

## Abstract

Species tree inference via summary methods that combine gene trees has become an increasingly common analysis in recent phylogenomic studies. This broad adoption has been partly due to the greater availability of genome-wide data and ample recognition that gene trees and species trees can differ due to biological processes such as gene duplication and gene loss. This increase has also been encouraged by the recent development of accurate and scalable summary methods, such as ASTRAL. However, most of these methods, including ASTRAL, can only handle single-copy gene trees and do not attempt to model gene duplication and gene loss. In this paper, we introduce a measure of quartet similarity between single-copy and multi-copy trees (accounting for orthology and paralogy relationships) that can be optimized via a scalable dynamic programming similar to the one used by ASTRAL. We then present a new quartet-based species tree inference method: ASTRAL-Pro (ASTRAL for PaRalogs and Orthologs). By studying its performance on an extensive collection of simulated datasets and on a real plant dataset, we show that ASTRAL-Pro is more accurate than alternative methods when gene trees differ from the species tree due to the simultaneous presence of gene duplication, gene loss, incomplete lineage sorting, and estimation errors.

## 1 Introduction

Evolutionary histories of genes and species can differ for several reasons [1], including incomplete lineage sorting (ILS), duplication and loss (duploss), gene transfer, and hybridization. Species tree inference is a central question in evolutionary biology and dealing with these sources of gene tree discordance is crucial. Many approaches have been proposed for species tree inference, including gene trees-species tree co-estimation [2–6] and species tree inference from sequence data [7–9]. However, the most scalable approach has remained a two-step process: first infer gene trees independently from sequence data and then combine them using a summary method. The goal of a summary method is to find the species tree best explaining the gene trees according to a model of gene tree discordance. While the ultimate goal is to develop summary methods modelling all sources of discordance, the literature mostly focused on separate causes.

A major family of summary methods focuses on duplication and loss processes producing multi-copy gene trees [10–15]. Most of these summary methods rely on maximum parsimony reconciliation [16] and aim at finding the species tree with the minimum reconciliation cost. Example methods include DupTree [10], its later extension iGTP [11, 12], DynaDup [15] and earlier similar dynamic programming algorithms [14]. Other methods take a more agnostic approach and minimize the distance between species trees and the gene trees without necessarily invoking specific reasons for discordance. Example methods of this type include MulRF [17] and *guenomu* [18]. However, a recent result asserts that the optimal solution to the optimization problem solved by MulRF is indeed a statistically consistent estimate of the species tree under a generic duplication-only model of gene evolution [19]. These methods are mostly designed to handle duplication and loss, and although in simulations some have reasonable accuracy under ILS and gene transfer [20], they have not been widely adopted by biologists.

Several summary methods target ILS as modelled by the multi-species coalescence (MSC) model [21, 22], and many of them are statistically consistent [e.g., 23–30]. However, the most successful summary method for ILS has arguably been ASTRAL [31], which, due to its high accuracy [32–34] and scalability [35, 36], has been used widely in biological analyses. ASTRAL, like several other methods [7, 24, 28], relies on dividing gene trees into unrooted four-taxon trees (called quartets), a feature that allows it to handle ILS and may contribute to its high accuracy. ASTRAL, however, was designed to handle single-copy gene trees reconstructed from sets of orthologous genes. This limitation has restrained the application scope of ASTRAL. For example, two studies on plant transcriptomes had to restrict species tree analyses with ASTRAL to the 400–800 putative single-copy gene trees, discarding thousands of available multi-copy genes [37, 38]. A surprising result asserts that treating gene copies as alleles of a same gene, a feature ASTRAL supports [39], is statistically consistent under a standard parametric model of gene duplication and loss and may be accurate [40]. Others have shown that random sampling of leaves works well empirically [41]. Beyond ASTRAL, several methods have focused on dividing multi-copy gene trees into single-copy genes without apparent duplications [42–46]. However, to our knowledge, no quartet-based methods *designed* to handle duplication and loss currently exist. Extending quartet-based methods to multi-copy gene trees is not trivial if we seek to correctly model orthology and paralogy.

Here, we introduce a quartet-based species tree inference method called ASTRAL-Pro (ASTRAL for PaRalogs and Orthologs). This method requires defining a measure of quartet similarity between single-copy and multi-copy trees accounting for orthology and paralogy. We define such a measure in a principled manner and show how to optimize it using dynamic programming. We test the method on an extensive collection of simulated datasets and on a real plant dataset.

## 2 Problem Definition

Let 𝒮 be a set of *n* species. Let us suppose that we are given a set of binary gene trees 𝒢, and, for each tree *G* ∈ 𝒢 with leaf set ℒ_*G*_ = {1 …*m*_*G*_}, we have a mapping *α*_*G*_: ℒ_*G*_ → 𝒮 specifying in which species each gene is sampled. For a rooted tree *G*, we denote the set of internal nodes in *G* by *I*(*G*), and, for each *u* ∈ *I*(*G*), we define ℒ_*G*_(*u*) as the set of leaves below *u*. We define two short-hands: *α*_*G*_(*A*) = {*α*_*G*_(*i*): *i* ∈ *A*} for *A* ⊂ ℒ_*G*_ and *α*_*G*_(*u*) = *α*_*G*_(ℒ_*G*_(*u*)) for a node *u*. The notation *G* ↾ *A* denotes *G* restricted to the set *A*.

We let Ω(*G*) be the multi-labelled tree obtained by replacing each leaf *i* ∈ ℒ_*G*_ with *α*_*G*_(*i*). Multiple copies of the same species in a gene tree *G* may be created by gene duplication. We assume that each duplication creates a new genomic locus (i.e., a position along the genome) and therefore, each locus, except the original one, has a parent locus (which may or may not have survived to the present day). Thus, each element of ℒ_*G*_ can be theoretically mapped to its parent locus, allowing us to “trace” the locus of each leaf to its ancestors.

In each gene tree *G*, we refer to a subset *Q* of four distinct elements of ℒ_*G*_ as a quartet. The subtree of a fully resolved tree *G* induced on a quartet *Q* exhibits two degree-three nodes. We refer to these nodes as *anchors of Q on G*. As shown in Fig. 1, for a rooted tree *G* and for a quartet *Q*, up to label permutations, *G* ↾ *Q* can only have two topologies: an *unbalanced* one (when one anchor descends from the other), denoted as *Q* ∠ *G*, and a *balanced* one (otherwise), denoted as *Q* ⊥ *G*. We say a tripartition (*P*_1_, *P*_2_, *P*_3_) of 𝒮 “can anchor” a quartet *Q* of *G* iff ∀_*i*_: *P*_*i*_ ∩ *α*_*G*_(*Q*) ≠ Ø.

**Figure 1:**
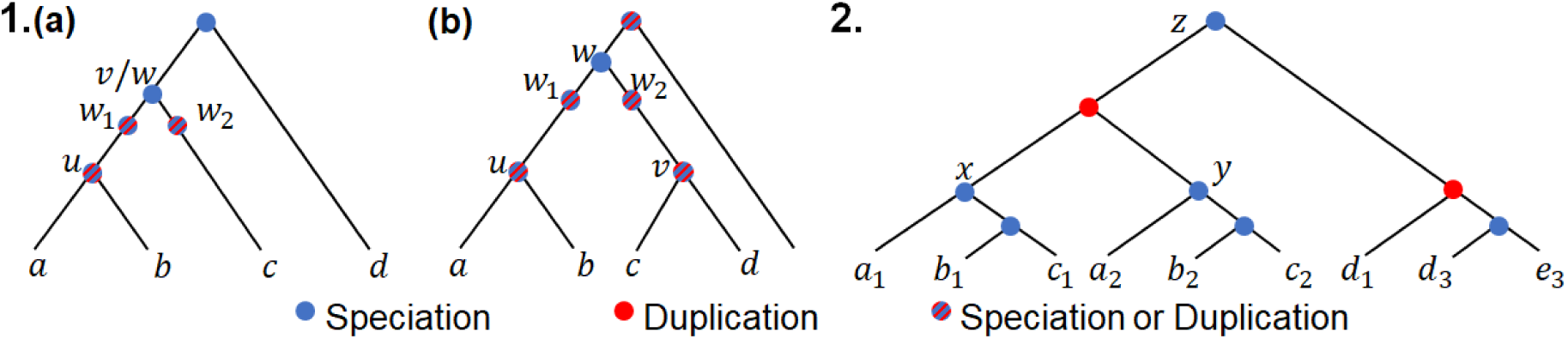
1. An example of a quartet *Q* = {*a, b, c, d*} with (*a*) unbalanced topology (*Q* ∠ *G*) and (*b*) balanced topology (*Q* ⊥ *G*). Anchors are *u* and *v*, and *w* = *ψ*_*G*_(*Q*) is the anchor LCA. While *w* has to be a speciation for *Q* to be considered a SQ, *u* and *v* can be either speciation or duplication. 2. An example of equivalent classes. Three equivalent classes are anchored on *z*: all eight quartets of the form {*a*_*i*_, *b*_*j*_, *d*_*k*_, *e*_3_}, of the form {*a*_*i*_, *c*_*j*_, *d*_*k*_, *e*_3_}, and of the form {*b*_*i*_, *c*_*j*_, *d*_*k*_, *e*_3_}, all with balanced topology. Anchored on *x*: two equivalent classes with unbalanced topology: {*a*_1_, *b*_1_, *c*_1_, *d*_1_} ∼ {*a*_1_, *b*_1_, *c*_1_, *d*_3_} and {*a*_1_, *b*_1_, *c*_1_, *e*_3_}. Anchored on *y*: two equivalent classes: {*a*_2_, *b*_2_, *c*_2_, *d*_1_} ∼ {*a*_2_, *b*_2_, *c*_2_, *d*_3_} and {*a*_2_, *b*_2_, *c*_2_, *e*_3_}.

### Definition 1

(Tagged trees). We say a rooted tree *G* is tagged if every internal node is tagged either as duplication or as speciation. A node *u* with children *u*_1_ and *u*_2_ can be a speciation if the three sets *α*_*G*_(*u*_1_), *α*_*G*_(*u*_2_), and *α*_*G*_(ℒ_*G*_ \ (ℒ_*G*_(*u*_1_) ∪ ℒ_*G*_(*u*_2_))) are mutually exclusive.

We note that these labels may or may not correspond to real speciation and duplication events. In particular, in the presence of deep coalescence across duplication events, a correct tagging corresponding to actual events may not be possible.

### Definition 2

(SQ). A quartet *Q* on a rooted tagged gene tree *G* is called a speciation-driven quartet (SQ) iff |*α*_*G*_(*Q*)| = 4 and the LCA of any three out of four leaves of *Q* is a speciation node. Equivalently, a quartet with topology *ab*|*cd* is a SQ if and only if its genes are all contained in different species and the LCA of either *a* or *b* with either *c* or *d* is tagged as speciation.

### Definition 3

(Quartet anchor LCA). Let *u* and *v* be anchors of a quartet *Q* on a *rooted* tree *G*. We refer to the LCA of *u* and *v* as the *anchor LCA* of *Q* on *G* and denote it as *ψ*_*G*_(*Q*).

The last definition is central to our approach. Note that anchors of a SQ can be speciations or duplications (Fig. 1) and thus SQs are not simply quartets with anchors being speciation nodes. Instead, they are quartets with a topology pre-determined by the speciation event represented by the anchor LCA, regardless of subsequent duplications and losses. Such subsequent duplications and losses may lead to multiple quartets originating from the same speciation event. Since these events include no new information on the speciation event, we count only SQs towards the quartet score of a species tree and weight them in a non-trivial way to avoid double-counting.

### Definition 4

(Equivalent SQs). Two SQs on the same 4 species are *equivalent* if they have the same anchor LCA; i.e., for two SQs, *Q*_1_ ∼ *Q*_2_ ⇔ *α*_*G*_(*Q*_1_) = *α*_*G*_(*Q*_2_) ∧ *ψ*_*G*_(*Q*_1_) = *ψ*_*G*_(*Q*_2_).

### Proposition 1.

*If Q*_1_ *and Q*_2_ *are equivalent SQs on G, then* Ω(*G* ↾ *Q*_1_) *and* Ω(*G* ↾ *Q*_2_) *are isomorphic*.

Thus, equivalent SQs have the same quartet topology when mapped to species. Proofs of all propositions and lemmas and sketches of proofs of all claims can be found in Appendix A.

Proposition 1 tells us that equivalent SQs do not provide any extra information with respect to each other, and therefore, it is reasonable to count all equivalent SQs as one unit when computing the quartet score of a species tree. This intuition is backed by the following proposition:

### Proposition 2.

*Assuming a correctly tagged tree G, for all equivalent SQs with a shared anchor LCA w, the three (in the unbalanced case) or four (in the balanced case) quartet leaves below w will all share an ancestral locus at the time of the speciation event corresponding to w*.

We can now provide a natural definition of the quartet score. The equivalence relation (Def. 4) partitions all quartets in equivalence classes and, by Proposition 1, for each equivalent class, we can define a unique quartet tree labelled by 𝒮. By Proposition 2, each class corresponds to an ancestral locus.

### Definition 5

(Per-locus (PL) Quartet Score). The per-locus quartet score of a species tree *S* with respect to a tagged gene tree *G* is the number of equivalent quartet classes with a quartet topology that matches the quartet tree induced by *S*. More formally,

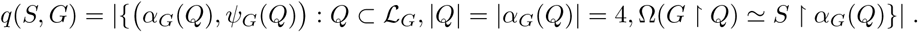

The PL quartet score of *S* with respect to a set of tagged gene trees 𝒢 is *q*(*S*, 𝒢) = ∑ _*G*∈ 𝒢_ *q*(*S, G*).

**Claim 1**. *If all nodes on the path between the root r and a node u are tagged as speciations, changing the root to any branch on the path does not alter the PL quartet score*.

### Problem 1

(Maximum per-Locus Quartet score Species Tree (MLQST) problem). *Given a set of rooted tagged gene trees* 𝒢, *find the species tree that maximizes the PL quartet score with respected to input gene trees, i*.*e*., arg max_*S*_ *q*(*S*, 𝒢).

## 3 Solving the MLQST problem using dynamic programming

We start by briefly describing the ASTRAL algorithm to solve a related problem (the MQSST problem), and then describe how we extend this approach to the MLQST problem.

### 3.1 Background: ASTRAL on single-copy gene trees

A node in a binary unrooted species tree forms a tripartition of 𝒮 that implies the topology for all quartets anchored at that node, allowing us to score it against 𝒢. Let *P* = *P*_1_|*P*_2_|*P*_3_ and *M* = *M*_1_|*M*_2_|*M*_3_ be two tripartitions, and let *I*_*ij*_ = |*M*_*i*_ ∩*P*_*j*_| Any species tree that displays *P* will share a certain number of quartets with any gene tree that displays *M*, and we call this number *QI*(*P, M*) (calculations below extends to multifurcations if *M* is a *d*-partition). Defining *G*_3_ as the set of all permutations of {1, 2, 3}, we have [31, 47]:

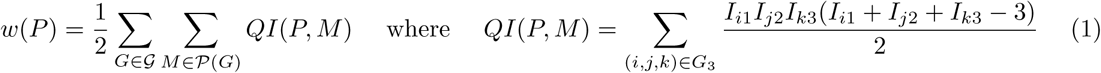

and 𝒫(*G*) is the set of partitions representing internal nodes of *G*. The quartet score of a species tree is simply the sum of the weights of its tripartitions. The division by half in *w*(*P*) is necessary because the sum counts each shared quartet twice (once at each anchor).

ASTRAL finds the tree *S* that maximizes the quartet score using dynamic programming. It recursively divides 𝒮 into subsets, in each step, choosing the division that maximizes the sum of the weights. To avoid exponential running time, instead of considering all ways of partitioning a set *A* ⊂ 𝒮 into *A*′ and *A* \ *A*′, we constrain the recursion to a given set of bipartitions. Let *X* be this set and *X*′ = {*A*: *A*|(𝒮 \ *A*) ∈ *X*} and *Y* = {(*C, D*): *C* ∈ *X*′, *D* ∈ *X*′, *C* ∩ *D* = Ø, *C* ∪ *D* ∈ *X*′}. Let *V* (*A*) be the quartet score of an optimal subtree on the cluster *A* and set *V* ({*a*}) = 0. Then,

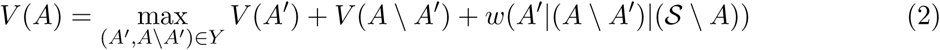

### 3.2 ASTRAL-Pro Algorithm

We extend here ASTRAL to multi-copy gene trees. The input to the new method, called ASTRAL-Pro (A-Pro for short), is a set of rooted tagged gene trees (see in Section 3.3 how unrooted gene trees can be rooted and tagged). This extension involves three changes. (*i*) To handle multi-copy gene trees, when computing the tripartition associated to each node, we use *α*_*G*_ to map labels to 𝒮. Here, instead of multi-sets, we create sets (counting multiple copies on each side only once). (*ii*) We change the weight calculation *w*(*P*) so that each equivalent class of quartets is counted once instead of twice, only at its LCA anchor. (*iii*) When computing *w*, we only sum over internal nodes tagged as speciations.

#### 3.2.1 Weight calculation

Let *w* be an internal node of *G* tagged as speciation and *P* = (*P*_1_|*P*_2_|*P*_3_) be a tripartition of 𝒮.

##### Definition 6.

We say that a SQ equivalent class with LCA anchor *w* in a gene tree *G* is mapped from left to a species tree tripartition *P* iff for each quartet *Q* in the equivalent class (*i*) *P* can anchor *Q* and (*ii*) the leaves *a* and *b* under the anchor of *Q* that appear first in a post-order traversal of *G* (e.g., *u* in Fig. 1) both map to the same side of *P* (that is, *α*_*G*_(*a*) ∈ *P*_*i*_, *α*_*G*_(*b*) ∈ *P*_*i*_ for some 1 ≤ *i* ≤ 3). We denote such quartets by 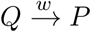.

We now state a set of lemmas, followed by the main result.

##### Lemma 1.

*If Q*_1_ ∼ *Q*_2_ *and* 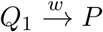, *then* 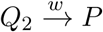.

##### Lemma 2.

*For a speciation node w with left child w*_1_ *and right child w*_2_, *let M*_1_ = *α*_*G*_(*w*_1_), *M*_2_ = *α*_*G*_(*w*_2_) *and M*_3_ = {*α*_*G*_(*z*): *z* ∈ ℒ_*G*_ \ ℒ_*G*_(*w*), *and LCA of w and z is tagged as speciation*}. *Let M*_*w*_ = (*M*_1_|*M*_2_|*M*_3_). *Recall I*_*ij*_ = |*M*_*i*_ ∩ *P*_*j*_|. *The number of SQ quartet equivalent classes anchored to w and mapped from left to the species partition P can be counted as follows:*

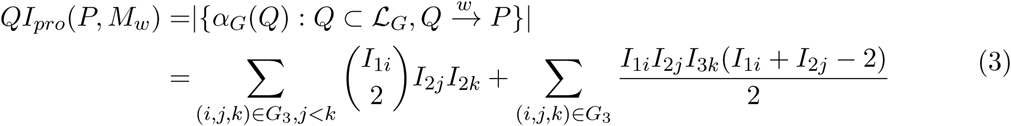

##### Lemma 3.

*If* Ω(*G* ↾ *Q*) ≃ *S* ↾ *α*_*G*_ (*Q*), *there exists a unique P* ∈ 𝒫 (*S*) *satisfying* 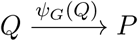.

##### Lemma 4.

*Let* **1**_*speciation*_(*w*) *be 1 for speciation nodes and 0 for duplication nodes and let*

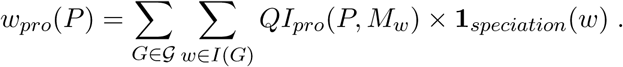

*Then: q*(*S*, 𝒢) =∑ _*P* ∈𝒫(*S*)_ *w*_*pro*_(*P*).

##### Theorem 1.

*The ASTRAL-Pro algorithm obtained by replacing w*(*P*) *function with w*_*pro*_(*P*) *in the ASTRAL dynamic programming solves the MLQST problem exactly if X* = 2^𝒮^.

*Proof*. By Lemma 4, arg max_*S*_ *q*(*S*, 𝒢) = arg max_*S*_ ∑ _*P* ∈𝒫(*S*)_ *w*_*pro*_(*P*). Thus, ASTRAL dynamic programming can solve the optimization problem exactly given the full search space (the argument is identical to that of ASTRAL and follows from the additive nature of *q*(*S*, 𝒢)). □

We now make two claims, and provide a sketch of proofs in Appendix A. Note that by Claim 3, ASTRAL-Pro has polynomial running time.

**Claim 2**. *For a set of gene trees* 𝒢 *including only speciations, the tree returned by ASTRAL-Pro is the same as the one returned by ASTRAL*.

**Claim3**. *The asymptotic running time of ASTRAL-Pro is O*(*D*|*X*|^1.73^) = *O*(*D*(*nN*)^1.73^) *where N* =∑_*G*∈𝒢_ |ℒ_*G*_| *and D denotes the number of unique gene tree tripartitions tagged as speciations*.

##### Algorithm 1 Gene tree tagging and rooting

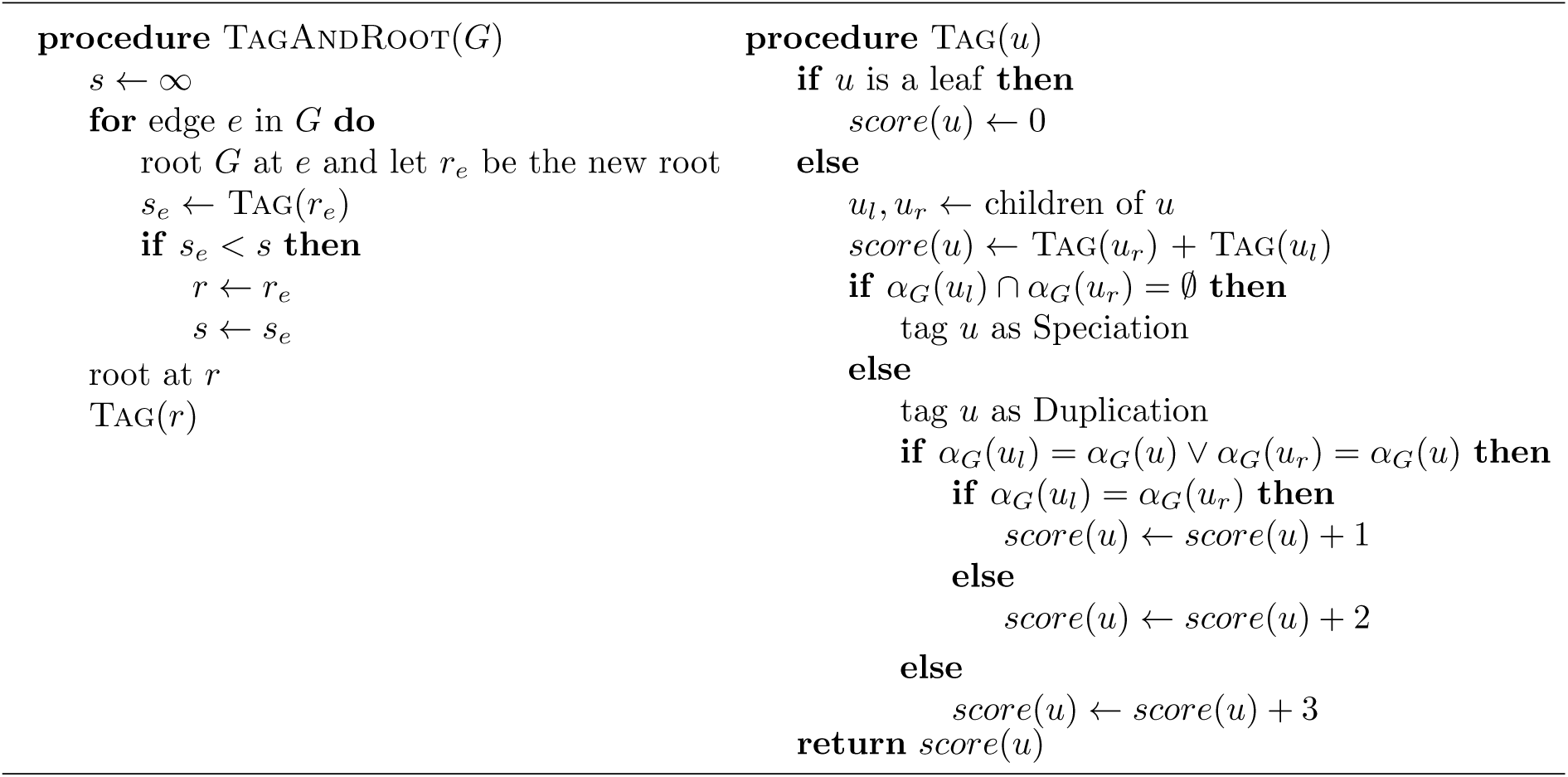

### 3.3 Tagging and rooting gene trees

Gene trees inferred from sequence data are neither rooted nor tagged. We use the heuristic Algorithm 1 to root and tag gene trees, noting that a partially-correct rooting suffices (Claim 1). Given a rooted tree, we tag a node as duplication *only if* the node cannot be tagged as speciation by Definition 1 (i.e., *observable duplication nodes* [43]); other nodes are *assumed* to be speciation.

For rooting, we seek the root position that minimizes the number of duplications and losses while allowing for free ILS. In each gene tree *G*, for two nodes *u* and *v* where *α*_*G*_(*u*) = *α*_*G*_(*v*), we explain all differences in topologies below *u* and *v* by invoking ILS (as opposed to duplication/loss). Then, three scenarios are possible for a node *u* with children *u*_*l*_ and *u*_*r*_. (*i*) When *u* is duplication and *α*_*G*_(*u*_*l*_) = *α*_*G*_(*u*_*l*_), we do not need to invoke any loss. One duplication suffices. (*ii*) If *α*_*G*_(*u*_*l*_) ⊂ *α*_*G*_(*u*_*r*_) or vice versa, we need one loss on *u*_*l*_ and an arbitrary amount of deep coalescence. (*iii*) Else, we need two losses (one in each side) and ILS to describe the differences. Algorithm 1 computes the number of duplication and loss events using this strategy, without penalizing ILS and fixing a cost of one for both duplications and losses. As described, it requires quadratic time per rooting and thus cubic to find optimal rooting. In our implementation, we used memoization to reduce this time to quadratic (details omitted). The LCA-based linear algorithm of Scornavacca *et al*. [43] could also be adapted.

#### 3.3.1 Search Space

We need to constrain the ASTRAL search space to bipartitions in a set *X*. To define *X*, we use a heuristic method relying on several strategies (see Algorithm 2). First, we use a sampling algorithm (SampleFull procedure) to create single-copy versions of each gene tree, creating a set ℱ. This sampling algorithm prunes the right (or left) subtrees below the highest duplication nodes in the tree, and recurses on each pruned tree, until no species has multiple copies. In addition, per gene, 2^*C*^ (default: *C* = 4) trees are sampled from ℱ, creating a multiset ℐ. This sampling can be probabilistic (taking each side of a duplication with probability 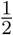) for high numbers of duplications. When the number of input trees is small, ℐ may become too small; in these cases, ℐ is augmented using another sampling algorithm (SampleExtra procedure). We provide ℐ as input to the algorithms implemented in ASTRAL-III for building the initial set *X* (as if -i ℐ is given to ASTRAL-III). Finally, we complete all trees from ℱ using the tree completion algorithm of ASTRAL-III and add the resulting bipartitions to *X*. All methods used guarantee that |*X*| grows polynomially with the number of species, gene trees, and duplication nodes.

##### Algorithm 2 Building set *X*. Default constant parameters: *C* = 4, *E*_*m*_ = 500, *E*_*s*_ = 4. The algorithm uses the (arbitrary) left/right orientation of children of a node as given in the input

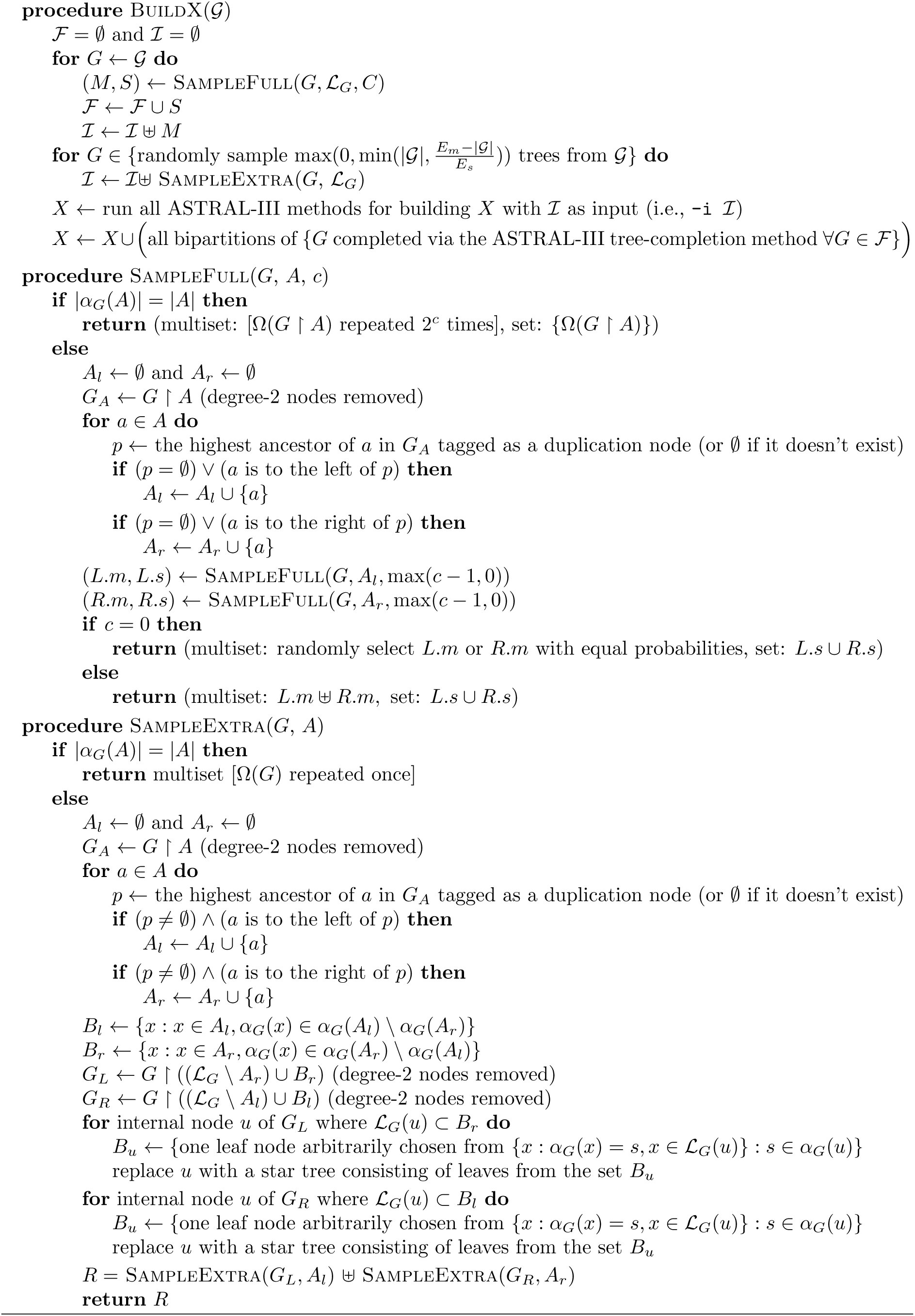

### 3.4 Statistical Consistency

When the input set 𝒢 has only speciation nodes, the MLQST problem reduces to the Maximum Quartet Support Species Tree (MQSST) problem solved by ASTRAL [31]. Thus, like the MQSST, the MLQST is NP-hard [48]. Moreover, the solution to MQSST problem is a statistically consistent estimator of the species tree under the MSC model and thus ASTRAL-Pro is also statistically consistent in absence of duplication.

In the presence of gene duplication and losses only, let us consider the birth-death model proposed by Arvestad *et al*. [49] and refer to it as the GDL model.

#### Proposition 3.

*Under the GDL model, every SQ in every correctly tagged rooted gene tree is isomorphic in topology to the species tree*.

Since all quartets in every equivalence class of SQs match the species tree, the per-locus quartet score will be maximized by the species tree. The following theorem follows.

#### Theorem 2.

*Under the GDL model [49], the solution to the MLQST problem is a statistically consistent estimator of the species tree given correctly rooted and tagged gene trees*.

In fact, we conjecture that ASTRAL-Pro is statistically consistent under the GDL model even when gene trees are imperfectly rooted and tagged. We leave the proof to future work. Finally, note that restricting to *X* does not impact statistical consistency, as each bipartition of the species tree has a non-zero chance of appearing in output of this algorithm.

## 4 Experiment setup

### 4.1 Datasets

We use new and existing simulated datasets as well as a biological dataset to test A-Pro.

#### 4.1.1 New simulated dataset (S25)

We perform a set of simulations using SimPhy [50] starting from a default model condition and adjusting five parameters (Table 1). We simulate 50 replicates per condition, and each replicate draws its parameters from prior distributions. Exact commands are given in Appendix B.

**Table 1:** Simulation settings for S25 dataset. *n*=number of ingroup species; *k*=number of genes; *τ* = tree height (generations); *λ*_+_= duplication rate; *λ*_−_= loss rate; *N*_*e*_= Haploid effective population size. Empirically, we estimate: *C* = mean number of copies per species minus one when *λ*_−_ = 0 and *n* = 25; ILS= mean RF distance between true gene trees and the species tree when *λ*_+_ = 0. MGTE = mean RF distance between true and estimated gene tree when *λ*_+_ = 0. See Table S1 for full parameters and Figures S1–S6 for full statistics.

|  |  |
| --- | --- |
| Default model | $n = 25; k = 1000; \tau \sim LN(21.25; 0.2); \lambda_+ = 4.9 \times 10^{-10}; \lambda_- = \lambda_+; N_e = 4.7 \times 10^8$<br>$C \approx 5; \text{ILS} \approx 70\%; \text{MGTE} = 15\% (500\text{bp}) \text{ or } 36\% (100\text{bp})$ |
| Controlling $\lambda_+, \lambda_-$<br>(duploss rate) | $\lambda_+ \in \{4.9, 2.7, 1.9, 0.52, 0\} \times 10^{-10} \lambda_- \in \{1, 0.5, 0.1, 0\} \times \lambda_+$<br>$C \approx \{5, 2, 1, 0.2, 0\}$ |
| Controlling $\lambda_+, N_e$<br>(dup rate, ILS) | $\{4.9, 1.9, 0\} \times 10^{-10}; N_e \in \{4.7 \times 10^8, 1.9 \times 10^8, 4.8 \times 10^7, 1.0 \times 10^4\}$<br>$\text{ILS} \approx \{70, 52, 20, 0\}\%; C \approx \{5, 1, 0\}$<br>$\text{MGTE} \approx \{15, 15, 15, 16\}\% (500\text{bp}) \text{ or } \approx \{36, 36, 36, 35\}\% (100\text{bp}) \text{ as } N_e \text{ changes}$ |
| Controlling $n$ | $n \in \{10, 25, 100, 250, 500\}$<br>$\text{MGTE} \approx \{15, 15, 17, 18, 18\} (500\text{bp}) \text{ or } \approx \{34, 36, 40, 43, 43\} (100\text{bp})$ |
| Controlling $k$ | $k \in \{25, 100, 250, 1000, 2500, 10000\}$ |

##### Default model

The species tree, simulated under the Yule process with birth rate 5 × 10^−9^ and the number of generations sampled from a log-normal distribution (mean 2.9 × 10^9^), has 25 ingroup and an outgroup species. Each replicate has 1000 true gene trees simulated under DLCoal with fixed haploid population size *N*_*e*_ = 4.7 × 10^8^. Gene trees have mean ILS level in [60%, 80%] range (mean 70%) across replicates (Fig. S2). The duplication rate *λ*_+_ = 4.9×10^−10^; when there is no loss, gene trees on average include 145 leaves (≈ 5 extra copies per species). The loss rate *λ*_−_ is set to *λ*_+_; with loss, gene trees have on average 43 leaves. The average number of duplication and loss events are 11 and 9, respectively, but variance is high (Fig. S1). For each gene, we use Indelible [51] to simulate gap-free nucleotide sequences along the gene trees using the GTR+Γ model [52] with 2 different sequence lengths: 500bp and 100bp. We then use FastTree2 [53] to estimate maximum likelihood gene trees under the GTR+Γ model. Gene tree estimation error, measured by the FN rate between the true gene trees and the estimated gene trees, depends on the sequence length and fluctuates significantly (from 0–100%) both within and across replicates (Fig. S3); mean error is 36% and 15% for 100bp and 500bp, respectively.

##### Controlling *λ*_+_, *λ*_−_

Here, we consider 5 × 4 = 20 conditions, changing duplication and loss rates. Our *λ*_+_ settings result in 0 to 5 extra copies per gene, and the 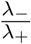 varies between 0 and 1 (Table 1; Fig. S4). All other parameters are identical to the default condition.

##### Controlling *λ*_+_, *N*_*e*_

Here, we consider 3 × 5 = 15 conditions, fixing *λ*_−_ to be equal to *λ*_+_, but changing *λ*_+_ and ILS levels (controlled by *N*_*e*_). Our *λ*_+_ settings result in 0 to 5 extra copies per gene, and the mean ILS level between true and estimated gene trees varies between 0 and 70% RF. (Table 1; Fig. S5) All other parameters are identical to the default model.

##### Controlling *n*

Fixing all parameters, we vary the number of ingroup taxa *n* from 10 to 500.

##### Controlling *k*

Fixing all parameters, we vary the number of gene trees *k* from 25 to 10,000.

#### 4.1.2 Existing simulations (S100)

We also used an existing dataset that Molloy and Warnow simulated [54] based on a real fungal dataset [55]. The simulation protocol of this dataset is similar to that of S25 dataset, with some notable differences. (*i*) The dataset included 100 species (no outgroup); species tree height, speciation rate, and mutation rates all differed from S25. (*ii*) Shorter gene alignments were also used, resulting in higher MGTE (25bp: 67%, 50bp: 52%, 100bp: 35%, 500bp: 19%). (*iii*) The duplication rate *λ*_+_ was set to 1×10^−10^, 2×10^−10^, or 5×10^−10^ (named 1, 2, and 5, respectively), and the duplication rate equaled the loss rate for all model conditions. (*iv*) ILS was much lower than S25; two conditions were simulated with *N*_*e*_ set to 1 × 10^7^ and 5 × 10^7^ (named 1 and 5, respectively), which result in 2% and 12% RF between true gene trees and the species tree. (*v*) Gene trees were estimated using RAxML instead of FastTree2.

#### 4.1.3 Biological data (1kp)

A transcriptome analysis of 103 plant species has been performed on 424 single-copy gene trees (out of thousands of genes) using both concatenation and ASTRAL [37]. In preliminary analyses, the authors had inferred multi-copy gene trees using RAxML from 9683 genes for 83 of those species, ranging in size between 5 and 2395 leaves. However, not being able to obtain an accurate species tree from the multi-copy gene trees, they abandoned the strategy in later analyses. The gene trees are available on Cyverse [56]. We used gene trees inferred from AA or first two codon positions (C12) as the original study.

### 4.2 Methods compared

We implemented A-Pro by extending ASTRAL-MP [36] and implementing Algorithms 1 and 2 as part of its native C++ library. We compare A-Pro to the following methods. Another method, STAG [57], is not included because of its poor performance on the S100 dataset [54], including that it fails to run on some model conditions (Fig. S8).

**DupTree** [10] infers a species tree from rooted or unrooted gene trees minimizing the duplication reconciliation cost [1] under the duplication-only model, but it does not model ILS. We provide DupTree with unrooted gene trees. We also tried iGTP, minimizing Dup-Loss score, but we only show results in supplement (Fig. S7) as it was almost universally worse than DupTree.

**MulRF** [17], based on an extension of the RF distance [58] to multi-labelled trees, is a hillclimbing method that aims at finding the tree with the minimum RF distance to the input. We use MulRF because of its advantage over other methods shown in previous studies [20].

**ASTRAL-multi** [39] is a feature of ASTRAL designed for handling multiple individuals. A recent paper (concurrently submitted) proposes to use ASTRAL-multi for multi-copy data [40]. Due to its high memory requirements, we were able to include it in only one experiment of S25.

## 5 Results

### 5.1 S25 dataset

#### 5.1.1 Controlling duplication and loss rates and the level of ILS

We start by experiments that change the duplication and loss rates (*λ*_+_, *λ*_−_) from the default condition (Fig. 2a). On true gene trees, A-Pro and DupTree are essentially tied in terms of accuracy, except for the case with no duplication and loss where A-Pro is perhaps slightly more accurate. Overall, the accuracy of A-Pro and DupTree is statistically indistinguishable under these conditions (*p*-value = 0.79 according to a multi-variate ANOVA test). Increasing *λ*_+_ *reduces* error (*p* < 10^−5^), perhaps because additional copies provide more information, akin to increasing the number of loci. Despite statistically significant increases (*p* = 0.006) in error as *λ*_−_ increases, both methods are quite robust to loss rates, losing at most 1.5% accuracy on average when *λ*_−_ = *λ*_+_ compared to no losses. MulRF has much higher error than other two methods, with errors that range between 10% and 17% across model conditions (we remind the reader that all these conditions have high ILS, a process that MulRF does not model).

**Figure 2:**
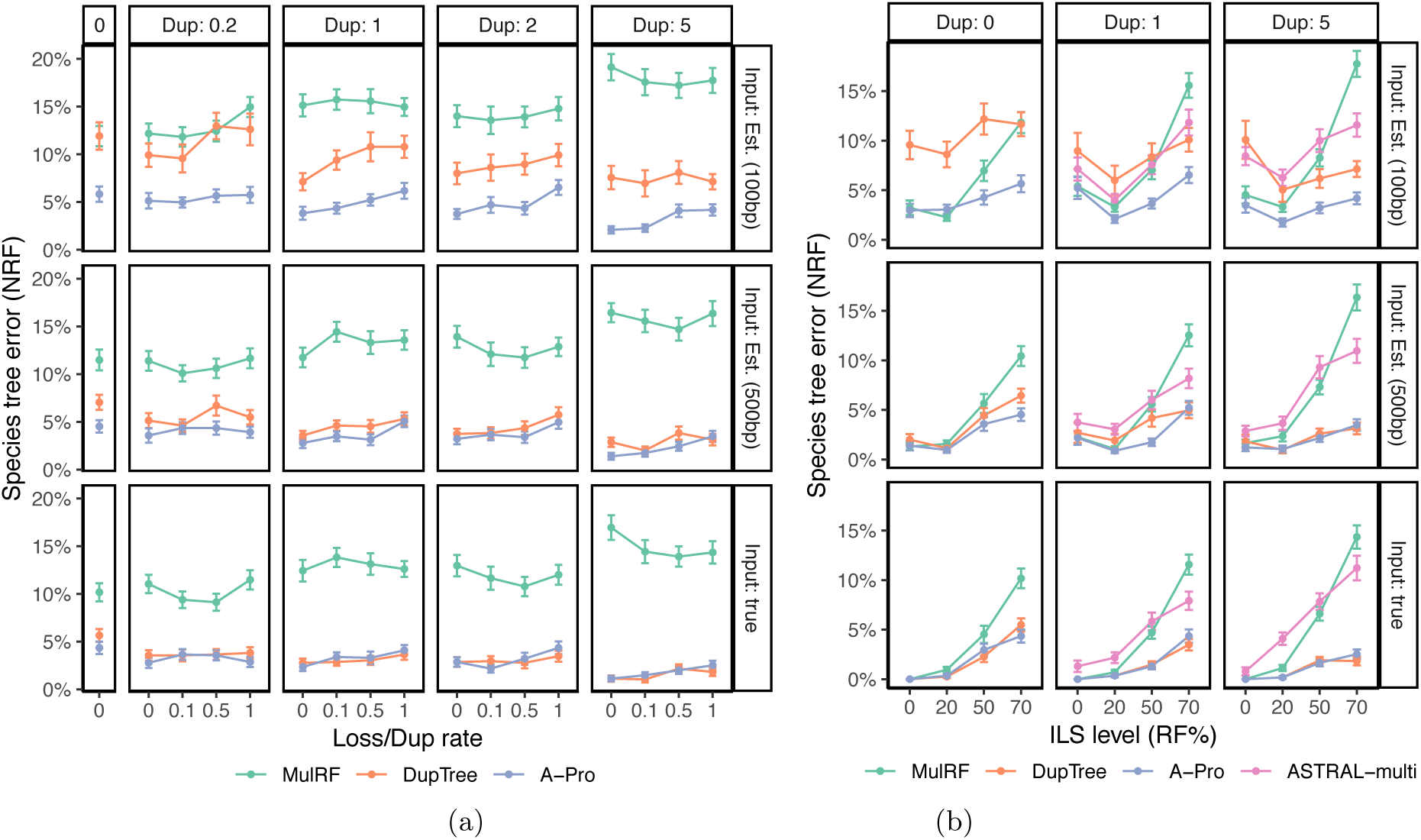
Species tree error on the S25 dataset for *n* = 25 ingroup species, *k* = 1000 gene trees, and both true and estimated gene trees from 100bp and 500bp alignments. (a) Controlling duplication rate (box columns; labelled by *C*) and the loss rate (x-axis; ratio of the loss rate to duplication rate). (b) Controlling the duplication rate (columns; labelled by *C*) and the ILS level (x-axis; NRF between true gene trees and the species tree for *λ*_+_ = 0). A-Pro and ASTRAL-multi are identical with *λ*_+_ = 0. See Table 1 for parameters and Fig. S7 for iGPT-duploss.

On estimated gene trees, the pattern changes, and the error of DupTree increases dramatically while A-Pro remains relatively accurate. When *λ*_+_ = *λ*_−_ = 0, DupTree has on average an 11.5% error whereas A-Pro has only a 4.5% error for 500bp. Adding duplications helps both methods but A-Pro remains more accurate. For example, with 100bp input gene trees, DupTree has an error between 50% to 260% higher than A-Pro. With low-error gene trees, differences are statistically significant (*p* < 10^−5^) but are more modest in magnitude (the error increase from DupTree to A-Pro across conditions by a median of 28%). The relative accuracy of A-Pro and DupTree is not a function of *λ*_−_ (*p* =0.8) but may depend on *λ*_+_ (*p* =0.05).

In terms of running time, on the default model condition, we observe that A-Pro is the fastest method, taking less than a minute on this dataset, followed closely by DupTree (Fig. S9). We will revisit running time of A-Pro on larger datasets below.

As we change the ILS level (Table 1), the reason for the poor performance of MulRF becomes clear (Fig. 2b). Without ILS, MulRF has excellent accuracy, often matching A-Pro and beating DupTree on low-error gene trees. As the ILS level increases (especially above 20%), the accuracy of MulRF deteriorates quickly. Overall, ILS has the strongest effect on accuracy (*p* ≪ 10^−5^) but its impact on methods vary (*p* ≪ 10^−5^). DupTree seems as tolerant of ILS as A-Pro, despite the fact that DupTree is not designed specifically for ILS, and both methods are much more tolerant of ILS than MulRF. Nevertheless, once again, DupTree shows extreme sensitivity to gene tree error. To summarize, DupTree is relatively tolerant of ILS but less tolerant of gene tree error; MulRF is tolerant of gene tree error but not of ILS; A-Pro is quite robust to both.

#### 5.1.2 Controlling the number of genes and species

Increasing the number of genes *k* in the most difficult case of high *λ*_+_, *λ*_−_, and ILS results in continued improvement in accuracy for A-Pro for every value we tested up to *k* = 10^4^ (Fig. 3a).

**Figure 3:**
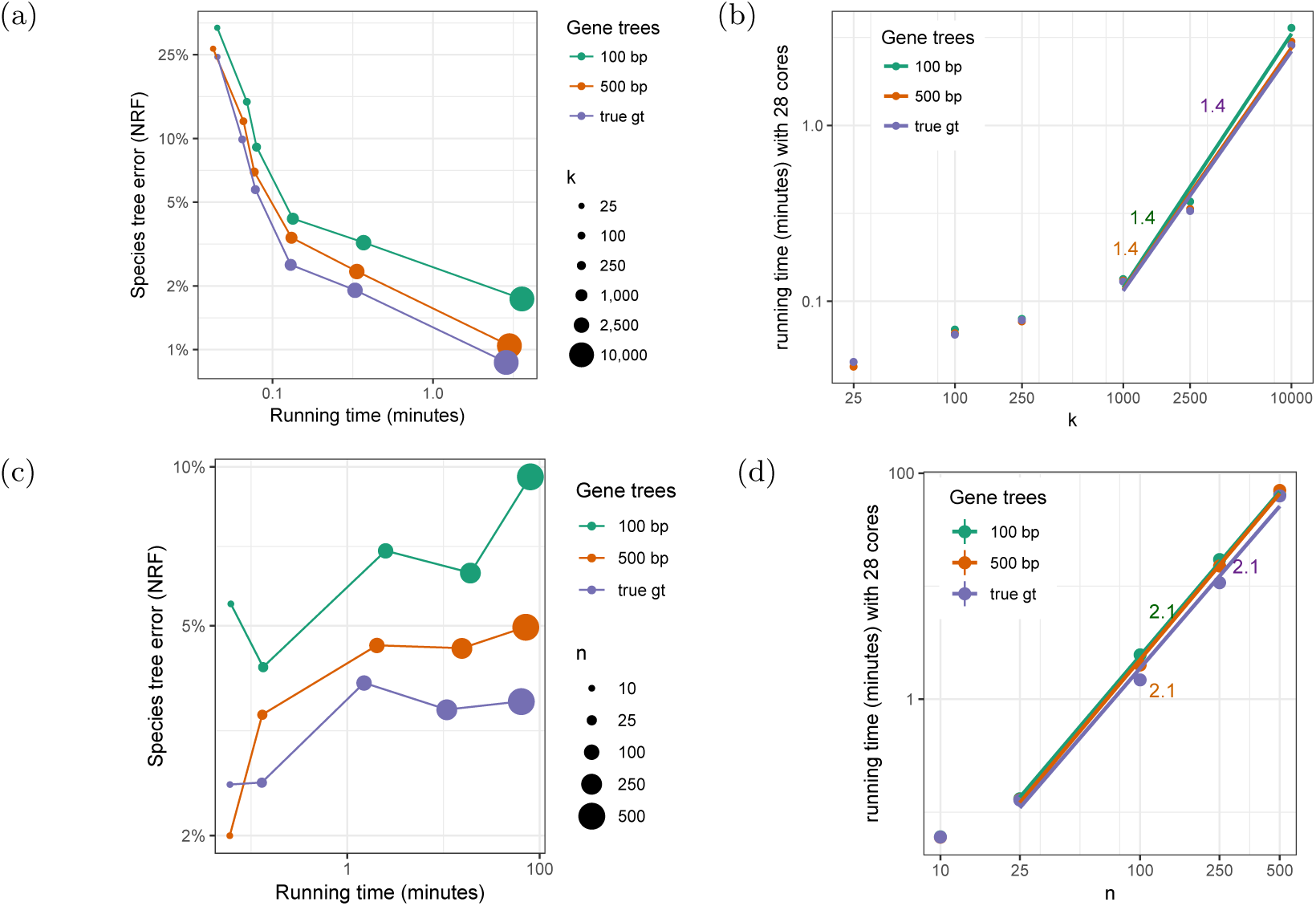
(a,c) Accuracy (y-axis) and running time (x-axis) of A-Pro as the number of genes *k* (a) or the number of species *n* (c) changes. Both axis are in log-scale. As *k* increases, accuracy increases. (b,d) The running time of A-Pro versus *k* (b) and *n* (d). We fit a line to the log-log plot of the running time only for *k* ≥ 1000 and *n* ≥ 25 as smaller runs are too fast to be reliable. We empirically estimate A-Pro to roughly proportionally with 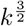 and *n*^*2*^.

With true gene trees, the error reduces from 26% with *k* = 25 to below 1% with *k* = 10^4^. Even with less accurate gene trees, the error reduces to below 2% with increased numbers of genes. Increasing *k* increases running time, which empirically grows with *k*^1.4^ (Fig. 3b). Nevertheless, using 28 cores, the running time was never more than 3.5 minutes even with *k* = 10^4^.

Increasing *n* from 25 to 500 shows that A-Pro is relatively robust to a large number of species (Fig. 3c). With true gene trees, the error ranges between 2.5% with 10 species to 3.5% with 500 species. With estimated gene trees, error ranges between 4.1% to 9.5% (for 100bp) and between 2% and 5% (for 500bp). Note that as *n* increases, the gene tree error also increases (Table 1; Fig S6). The running time of A-Pro increases roughly quadratically with *n* (Fig. 3d) but is below 2 hours (given 28 cores) even for *n* = 500 (recall that *k* = 1000).

### 5.2 S100 dataset

Patterns of performance on the S100 dataset are consistent with the S25 dataset (Fig. 4). DupTree is highly accurate with true gene trees and gene trees with low estimation error but quickly degrades in accuracy as gene tree error increases. MulRF is less sensitive to gene tree error but is more sensitive to the ILS level (which is always moderate or low on this dataset). As in S25, here, we see that using ASTRAL-multi to handle duplication and loss is not beneficial. A-Pro works the best overall, ranking first in terms of mean error (rounded to two significant digits) in 105 out of 120 test conditions (Table S2). The second best method is MulRF on this dataset, which is not surprising given the low ILS levels in this dataset.

**Figure 4:**
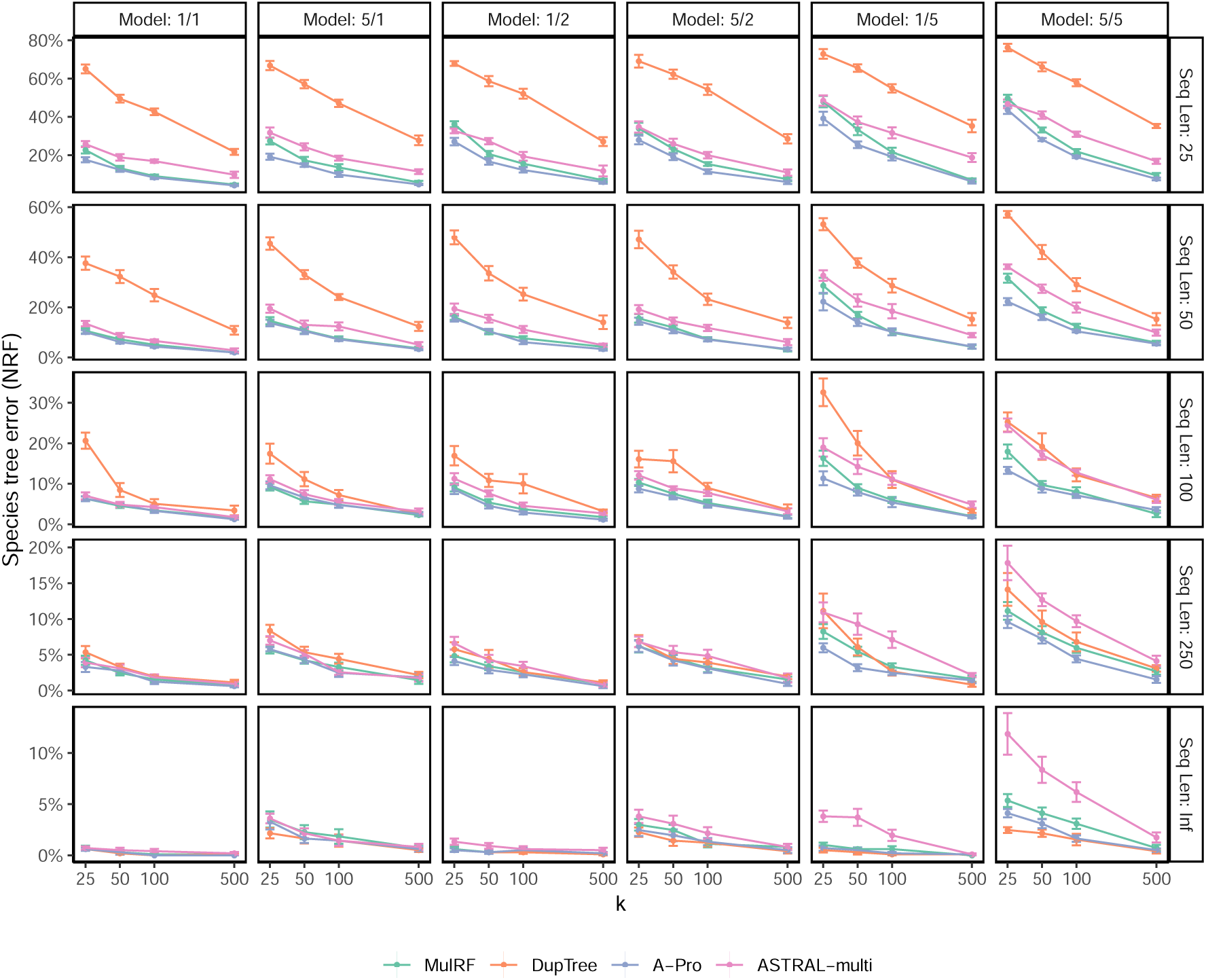
Species tree error on S100 dataset. We compare the species tree error of the four methods, showing mean and standard error over 10 replicates for each model condition, with varying numbers of genes (*k*) and sequence lengths (with Inf signifying true gene trees). Model conditions are labeled as *a/b* where *a* is the level of ILS (1 or 5) and *b* is the duplication/loss rate (1, 2, or 5).

### 5.3 Biological plant dataset

On the plant dataset [37], A-Pro on AA gene trees returned a species tree (Fig. 5) that was largely similar to the main single-copy ASTRAL tree reported by the original study (only 4 differences). Using C12 gene trees resulted in only one change, swapping Chara and Coleochaetales. In contrast, DupTree differed from the ASTRAL tree in 33 out of 77 branches (21/77 for iGTP-DupLoss) and violated many known biological relationships (Fig. S10). The A-Pro trees are consistent with ASTRAL for major groups, including placing Zygnematales (not Chara) as sister to all land plants, the placement of Amboerlla as sister to the rest of angiosperms, and monophyly of Bryophytes (liverworts, mosses, and hornworts).

**Figure 5:**
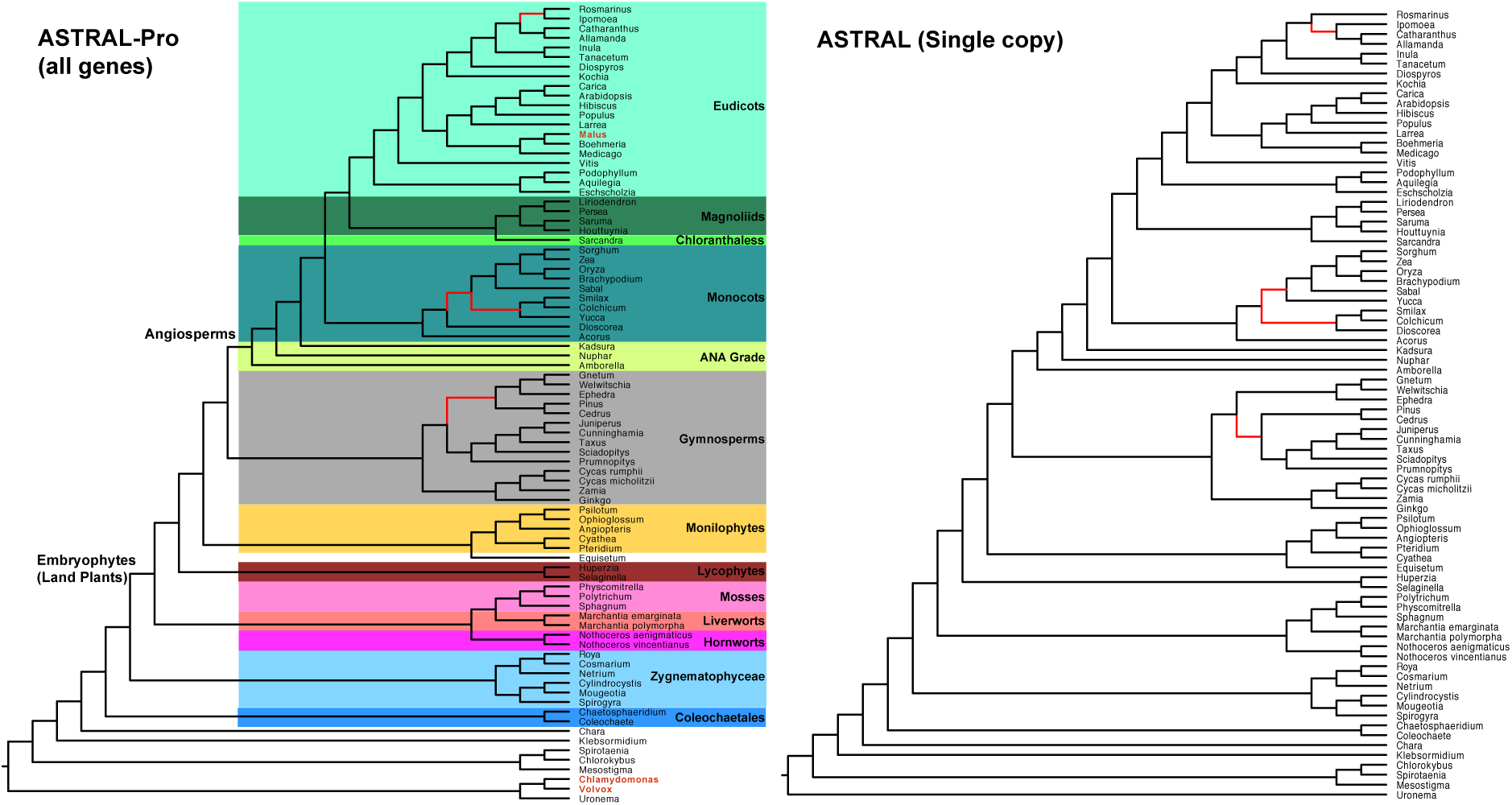
ASTRAL and ASTRAL-Pro on biological plant paper. ASTRAL-Pro is run on 9683 multi-copy AA gene trees available online [56]. ASTRAL is run on 424 single-copy gene trees and was reported previously [37]. Three genomes, shown in red, were present in multi-copy gene trees but not in the single-copy analyses. The single-copy tree includes 23 extra species that were not in the multi-copy data and are removed here. The two trees are very similar and differ in only four branches shown in red.

The four changes between the ASTRAL and A-Pro trees are interesting. In A-Pro, unlike ASTRAL, Rosmarinus and Ipomoea are grouped together, which is likely the correct result as these species are in the same order (Lamiales). The position of genus Yucca in the A-Pro tree has changed; interestingly, a recent update to this paper using > 1, 000 species [38] (which samples close genera Asparagales and Liliales) finds Yucca in a position identical to A-Pro. Thus, the A-Pro placement is more likely to be correct. Most consequentially, A-Pro, unlike ASTRAL, recovers the GnePine hypothesis combining Gnetales and Pinaceae, a hypothesis suggested by several studies [59–62] and all concatenation analyses of 1kp [37]. Interestingly, the new 1kp paper [38] uses DiscoVista [63] to examine quartet frequencies around this branch and detects that the second and third most frequent quartets do not match (0.4 vs. 0.1) and are heavily skewed towards GnePine, making the resolution obtained in ASTRAL less reliable.

## 6 Discussions

We developed a “per-locus” quartet-based measure of similarity between multi-copy gene trees and a species tree. The measure relies on internal nodes of gene trees being tagged as speciation or duplication. Somewhat counter-intuitively, despite being a quartet measure, it needs *partially* rooted trees (Claim 1). The measure defines an equivalence relationship on quartets and counts each equivalent class only once, avoiding double-counting quartets that are bound to have identical topologies. Avoiding double-counting is at the heart of the approach and likely is a main reason behind its high accuracy on simulated and empirical data we tested.

Astral-Pro, which maximizes the per-locus quartet score, is statistically consistent under MSC and GDL models. This makes one hope that it may also be consistent under both causes of discordance combined. The DLCoal model [55] accounts for ILS, duplication, and loss. Under this model, each duplication immediately creates a daughter locus, which is unlinked from the parent locus; the duplication event gets fixed in all species. Gene trees are seen as generated by first producing a locus tree via a birth-death process that runs on the species tree and then running a MSC process on the locus tree. Because the loci are considered as unlinked, the coalescence processes occur independently between the parent and daughter loci (but the daughter MSC process is “bounded” at the time of duplication). Due to the independence of loci, dividing a multi-copy gene family into its constituent loci can give us distributions on gene tree topologies that behave similarly (though not identically) to the MSC model. The per-locus metric *seeks* to count quartet topologies across loci as they existed at the time of speciation events relevant to a quartet (i.e., at the time of the anchor LCA). When successful, it counts only topologies that are drawn from independent coalescent processes. However, complicated scenarios involving a combination of duplications, losses and ILS can lead to incorrectly tagged gene trees. These scenarios create complications that need to be addressed. We leave it to the future work to study whether ASTRAL-Pro is statistically consistent under the DLCoal model.

To get rooted and tagged gene trees, we used the maximum parsimony principle, with duplication and loss each penalized equally and deep coalescence not penalized at all. There is a large literature on various ways of tagging and rooting gene trees [e.g., 64–66], including other penalties for the duplication and loss events (e.g., there is a suggestion of losses having half the penalty of duplications [67]). It may also be possible to improve tagging of gene trees using probabilistic orthology inference [68, 69] or using synteny information [70, 71]. However, these methods often require a species tree. It may possible to use A-pro in an iterative fashion, where the species tree is inferred, gene trees are re-tagged and re-rooted, and a new species tree is inferred Future work should explore these approaches.

Quartet-based methods for handling multi-copy gene trees are not abundant. Besides our method, one can attempt to sample single-copy gene trees [41], an approach that we plan to test in the future. Most recently, there has been theoretical and empirical evidence that simply treating gene copies as alleles may be sufficient [40]. We showed that this alternative, although attractive in theory, is less accurate and less scalable than A-Pro. We are unaware of other quartet-based species tree inference methods for multi-copy input.

A-Pro has some limitations. Most importantly, in its current form, it can only handle binary trees, which reduces its ability to handle gene tree error [47]. While A-Pro is more robust to gene tree error than alternatives, combining it with co-estimation [3] and tree fixing [72–77] may further improve its accuracy. Future work should also explore ways to extend A-Pro so that it can handle polytomies in input gene trees. Finally, with more algorithmic development, it should be possible to provide all the features ASTRAL provides, including branch length and Local-PP [78], polytomy test [79], and visualization of discordance [63]. All these features should be adopted to A-Pro in the future.

## Data availability

The code is available at https://github.com/chaoszhang/A-pro and data are made available at https://github.com/chaoszhang/duploss-pipeline.git.

## Acknowledgments

S.M and C.Z were supported by the National Science Foundation (NSF) grant III-1845967. E.K.M. was supported by the Ira and Debra Cohen Graduate Fellowship in Computer Science.

## A Proofs

### Proof of Proposition 1.

Denote *Q*_1_ = {*a, b, c, d*} and 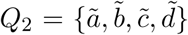 (with obvious correspondence of labels). Let *w* be the anchor LCA and note that anchor LCA is the LCA of three (if *Q*_1_ ∠ *G*) or four (if *Q*_1_ ⊥ *G*) of the quartet leaves; thus, by Definition 2, *w* is a speciation node or otherwise *Q*_1_ would not be a SQ. Let denote by *w*_1_ and *w*_2_ the children of *w*; by Definition 1, *α*_*G*_(*w*_1_), *α*_*G*_(*w*_2_) and the remaining leaves (*U* = *α*_*G*_(ℒ_*G*_ − ℒ_*G*_(*w*_1_) − ℒ_*G*_(*w*_2_))) must be mutually exclusive. In the unbalanced case, given that *a, b* ∈ ℒ_*G*_(*w*_1_), *c* ∈ ℒ_*G*_(*w*_2_), *d* ∈ *U*, mutual exclusivity is possible only if 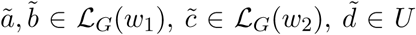. In the case of balanced topology, mutual exclusivity of *α*_*G*_(*w*_1_) and *α*_*G*_(*w*_2_) and the fact that *a, b* ∈ ℒ_*G*_(*w*_1_) and *c, d* ∈ ℒ_*G*_(*w*_2_) implies that 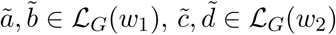. Thus, in either case, Ω(*G* ↾ *Q*_1_) ≃ Ω(*G* ↾ *Q*_2_). □

### Proof of Proposition 2.

Each node of a gene tree represents an ancestral or present-day gene and thus belongs to a locus. The children of a speciation node stay in the same locus that their parent, while for a duplication node we have that exactly one of the two children change locus and the other stays in the same locus than its parent. Therefore, all nodes under *w*, which is a speciation node, belong to the descendants (including itself) of the locus to which *w* belongs, and when tracing back to the time of speciation event *w*, they will lead to the same locus. Since all equivalent classes share the same anchor LCA, the result follows. □

### Proof of Lemma 1.

Note that *P* can anchor *Q*_1_ only iff any species tree that includes *P* must match the gene tree topology for *Q*_1_ By Proposition 1, due to equivalence of *Q*_1_ and *Q*_2_, we infer *Q*_2_ must (i) match the same species quartet set as *Q*_1_ and (ii) share the same anchor LCA *w*. Thus, *P* can also anchor *Q*_2_. (iii) When *Q*_1_ ∠ *G* as shown in Figure 1, 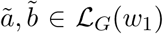 are the leaves mapped to the quartet tree and thus mapped to the same partition as *a, b*; similarly, when *Q*_1_ ⊥ *G*, the pair of leaves under left subtree of the anchor LCA of both quartets map to the same partition of *P*. □

### Proof of Lemma 2.

First note that:

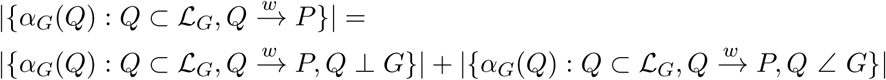

We compute each part separately. Recall here that, since 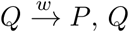, is a SQ quartet and thus |*Q*| = |*α*_*G*_(*Q*)| = 4.

When *Q* ⊥ *G*, let *Q* = {*a, b, c, d*} with *α*_*G*_(*a*), *α*_*G*_(*b*) ∈ *M*_1_, *α*_*G*_(*c*), *α*_*G*_(*d*) ∈ *M*_2_. Since 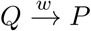, leaves *α*_*G*_(*a*) and *α*_*G*_(*b*) must be in the same partition of *P*. When *α*_*G*_(*a*), *α*_*G*_(*b*) ∈ *P*_1_, leaves *α*_*G*_(*c*) and *α*_*G*_(*d*) must be in partition *P*_2_ and *P*_3_ respectively since *P* can anchor *Q*. W.l.o.g., we can assume *α*_*G*_(*c*) ∈ *P*_2_. Therefore, *α*_*G*_(*a*), *α*_*G*_(*b*) ∈ *M*_1_ ∩ *P*_1_, *α*_*G*_(*c*) ∈*M*_2_ ∩ *P*_2_, *α*_*G*_(*d*) ∈ *M*_2_ ∩ *P*_3_. The number of such *α*_*G*_(*Q*) is 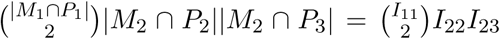. Similarly when *α*_*G*_(*a*), *α*_*G*_(*b*) ∈ *M*_1_ ∩ *P*_2_ and *α*_*G*_(*a*), *α*_*G*_(*b*) ∈ *M*_1_ ∩ *P*_3_, the number of such *α*_*G*_(*Q*) is 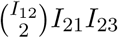 and 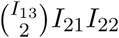 respectively. Thus,

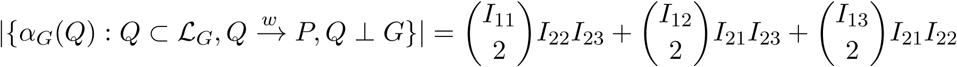

Similarly, when *Q* ∠ *G*, let *Q* = {*a, b, c, d*} with *α*_*G*_(*a*) and *α*_*G*_(*b*) in the same partition of *P*. Notice that, in the unbalanced case, *α*_*G*_(*a*) and *α*_*G*_(*b*) can be both either in *M*_1_ or either in *M*_2_, and since *c* and *d* are not interchangeable as in the balanced case, we can have *α*_*G*_(*a*), *α*_*G*_(*b*) ∈ *P*_*i*_, *α*_*G*_(*c*) ∈ *P*_*j*_, *α*_*G*_(*d*) ∈ *P*_*k*_ for (*i, j, k*) with any permutation of (1, 2, 3), from the definition of *P* anchoring *Q*. All together we have 12 cases.

In the case that *α*_*G*_(*a*), *α*_*G*_(*b*) ∈ *P*_1_, *α*_*G*_(*c*) ∈ *P*_2_, *α*_*G*_(*d*) ∈ *P*_3_, and *α*_*G*_(*a*), *α*_*G*_(*b*) ∈ *M*_1_, we have *α*_*G*_(*a*), *α*_*G*_(*b*) ∈ *M*_1_ ∩ *P*_1_, *α*_*G*_(*c*) ∈ *M*_2_ ∩ *P*_2_, and *α*_*G*_(*d*) ∈ *M*_3_ ∩ *P*_3_. The number of such *α*_*G*_(*Q*) is 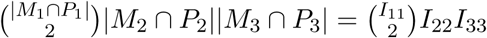. The other 11 permutations are similar. In total,

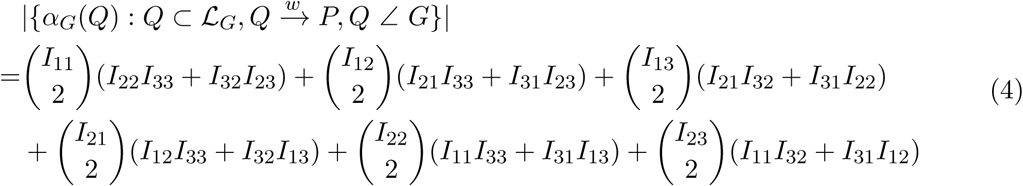

Thus,

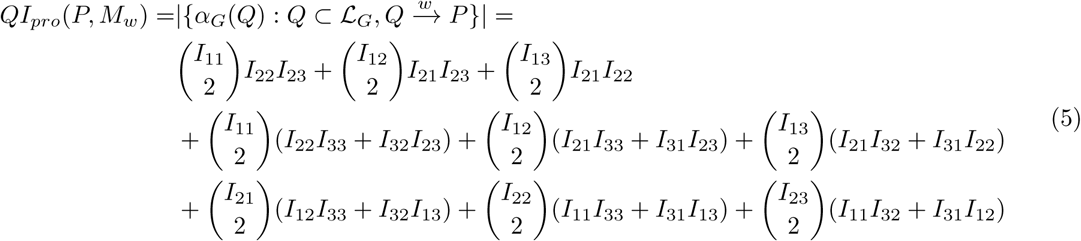

With simple manipulations, it can be shown that the right hand side of this equation can be rewritten as:

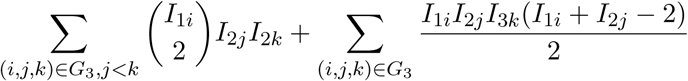

□

### Proof of Lemma 3.

Let Ω(*G* ↾ *Q*) be designated by *ab*|*cd* and assume w.l.o.g that the anchor corresponding to *a* and *b* is the first anchor observed on the post-order traverse of *G*. It is easy to show (see [31]) that if Ω(*G* ↾ *Q*) ≃*S* ↾ *α*_*G*_(*Q*) there exist exactly two tripartitions *P* ^1^ and *P* ^2^ in P(*S*) that imply a quartet topology that matches Ω(*G* ↾ *Q*) (condition (*ii*) of Definition 6). Each of the two tripartitions has two leaves of *α*_*G*_(*Q*) in one of its parts and the other two leaves fall on two different parts. Also, the two leaves that are together can only be *a* and *b* or *c* and *d* and thus, only one of *P* ^1^ and *P* ^2^ would include both *a* and *b* in the same part. Therefore, by condition (*iii*) of Definition 6, exactly one of 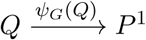 and 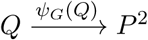 can be true. □

### Proof of Lemma 4.

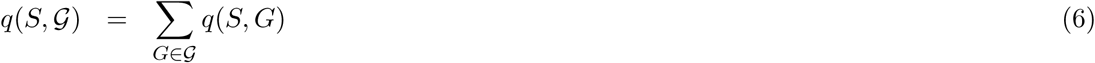

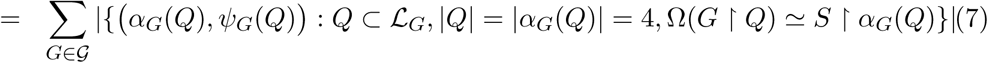

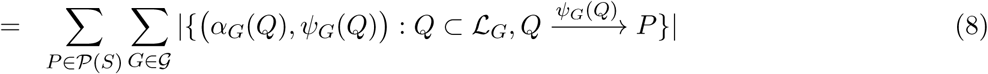

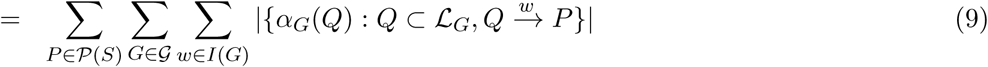

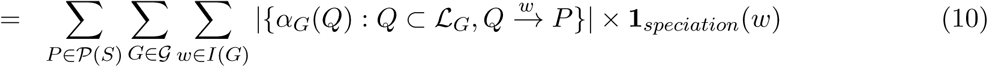

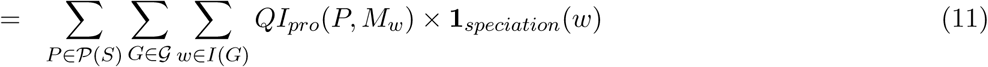

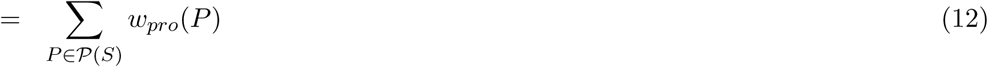

The first two lines are implied by Definition 5. Equation (8) follows from Lemma 1 and Lemma 3 that together establish that each equivalence class of quartets maps to exactly one *P*. Equation (9) follows from Definition 4 combined with a simple rearrangement obtained by counting unique tuples once. Equation (10) follows from the fact that when *w* is a duplication node, 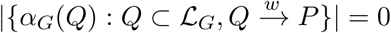. Equation (11) follows from Lemma 2. □

*Proof sketch of Claim 1*. The rooting that minimizes the number of duplications and losses (#duploss for short) in Alg. 1 may not be unique. In particular, if a rooted tree *G* minimizes #duploss, then rooting it at any branch such that the path between the parent node of the branch and the current root (including the two end nodes) does not contain any duplication node will also minimize #duploss. We call a correctly-tagged gene tree partially-correctly-rooted if the path between the parent node of the branch where it is rooted and the root in the correctly-rooted tree does not contain any duplication node. In particular, when gene trees do not have duplications, then any rooting of a gene tree is partially-correctly. We observe that the equivalent classes of quartets in all partially-correctly-rooted trees stay the same (although *all* quartet trees in the same equivalent class may change from balanced to unbalanced or vice versa), and thus any partially-correct-rooting of gene trees will result in the same species tree. □

*Sketch of proof of Claim 2*. When G only includes speciation nodes, regardless of rooting, each quartet is a SQ. Since each leaf corresponds to distinct taxa in the species tree, each quartet equivalent class contains only one quartet. Therefore, each quartet is counted exactly once and thus ∑_*P* ∈𝒫 (*S*)_ *w*_*pro*_(*P*) = ∑_*P* ∈𝒫 (*S*)_ *w*(*P*) regardless of rooting. □

*Sketch of proof of Claim 3 (Running time of ASTRAL-Pro)*. Let

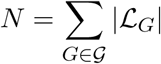

denote the sum of the number of leaves in the gene trees. Then the number of anchor LCAs in all gene trees is *O*(*N*). Let *D* denote the number of unique gene tree tripartitions tagged as speciations and note *D* = *O*(*N*). By only counting each unique gene tree tripartition once against each species tree tripartition, the running time of ASTRAL-Pro becomes *O*(*D*|*X*|^1.73^) (by an argument that is identical to that provided for ASTRAL-III [31] and follows from results of Kane and Tao [80]). However, while ASTRAL-III guarantees |*X*| = *O*(*nk*) with *k* = |𝒢|, in ASTRAL-Pro, in the presence of duplications, |*X*| can be large; in particular with our sampling algorithm (Alg. 2), |*X*| = *O*(*nN*). Thus, the running time of A-Pro is *O*(*D*(*nN*)^1.73^). Note that this analysis is not tight and can be made more precise in the future. Also, in the future, we will explore sub-sampling a constant number of trees from the output of Alg. 2 per gene tree, which will limit the |*X*| = *nk* and thus limit the running time of ASTRAL-pro to *O*(*D*(*nk*)^1.73^). □

### Proof of Proposition 3.

Under GDL, besides leaves, each internal node *u*_*G*_ ∈ *I*(*G*) in a gene tree *G* corresponds to an internal node *u*_*S*_ ∈ *I*(*S*); if *u*_*G*_ is a duplication node, *u*_*S*_ is the node down the branch in *S* where the duplication event happened, and if *u*_*G*_ is a speciation node, *u*_*S*_ is the respective speciation node. It is easy to see that *α*_*G*_(*u*_*G*_) ⊂ ℒ_*S*_(*u*_*S*_). For each SQ quartet *Q* = {*a, b, c, d*}, assuming w.o.l.g that *G* ↾ *Q* has unrooted topology *ab*|*cd*, let *w*_*G*_ = *ψ*_*G*_(*Q*), and *u*_*G*_ and *v*_*G*_ be the children of *w*_*G*_. Let *u*_*G*_, *v*_*G*_, and *w*_*G*_ correspond to *u*_*S*_, *v*_*S*_, and *w*_*S*_ in *S*, respectively. Since *w*_*G*_ is a correctly tagged speciation node, *u*_*S*_ and *v*_*S*_ are descendants from different children of *w*_*S*_.

When *Q* ⊥ *G*, assuming w.o.l.g. *a, b* ∈ ℒ_*G*_(*u*_*G*_) and *c, d* ∈ ℒ_*G*_(*v*_*G*_), we get *α*_*G*_(*a*), *α*_*G*_(*b*) ∈ ℒ_*S*_(*u*_*S*_) and *α*_*G*_(*c*), *α*_*G*_(*d*) ∈ ℒ_*S*_(*v*_*S*_) and thus *α*_*G*_(*a*)*α*_*G*_(*b*)|*α*_*G*_(*c*)*α*_*G*_(*d*) is induced by *S*.

When *Q* ∠ *G*, assuming w.o.l.g. *a, b* ∈ ℒ_*G*_(*u*_*G*_), *c* ∈ ℒ_*G*_(*v*_*G*_), and *d* ∉ ℒ_*G*_(*w*_*G*_), we get *α*_*G*_(*a*), *α*_*G*_(*b*) ∈ ℒ_*S*_(*u*_*S*_) and *α*_*G*_(*c*) ∈ ℒ_*S*_(*v*_*S*_). Since *d* is not under *w*_*G*_, *α*_*G*_(*d*) and *w*_*S*_ are under different children of the species tree node to which the LCA of *d* and *w*_*G*_ corresponds. Therefore, *α*_*G*_(*d*) ∉ ℒ_*S*_(*w*_*S*_) and thus *α*_*G*_(*d*) ∉ ℒ_*S*_(*u*_*S*_); since *α*_*G*_(*a*) ∈ ℒ_*S*_(*u*_*S*_) and *α*_*G*_(*b*) ∈ ℒ_*S*_(*u*_*S*_), it follows that *α*_*G*_(*a*)*α*_*G*_(*b*)|*α*_*G*_(*c*)*α*_*G*_(*d*) in *S*. □

## B Simulation details

Simphy command for default parameters:

simphy –s l f: 2 5 –rs 50 –r l f: 1000 –rg 1 –sb f: 0. 0 0 0 0 0 0 0 0 5 –sd f: 0 –s t ln: 2 1. 2 5, 0. 2 –so f: 1 –s i f: 1 –sp f: 470000000 –su ln: – 21. 9, 0. 1 –hh f: 1 –hs ln: 1. 5, 1 –hl ln: 1. 5 5 1 5 3 3, 0. 6 9 3 1 4 7 2 –hg ln: 1. 5, 1 –cs 9644 –v 3 –o d e f a u l t –ot 0 –op 1 –lb f: 0. 00000000049 –ld f: 0. 00000000049 –l t f: 0

Other settings use a similar command with parameters changed according to the table below.

**Table S1:** Simphy parameters for all experiments.

| Parameter name | Parameter value |
| --- | --- |
| Default Parameters |  |
| Speciation rate | 5e-9 |
| Extinction rate | 0 |
| Locus trees | 1000 |
| Gene trees | 1 |
| Number of leaves | 25 + an outgroup |
| Ingroup divergence to the ingroup ratio | 1.0 |
| Generations | LogN(21.25,0.2) |
| Haploid effective population size | 4.7e+8 |
| Global substitution rate | LogN(-21.9,0.1) |
| Lineage specific rate gamma shape | LogN(1.5,1) |
| Gene family specific rate gamma shape | LogN(1.551533,0.6931472) |
| Gene tree branch specific rate gamma shape | LogN(1.5,1) |
| Duplication rate | 4.9e-10 |
| Loss rate to duplication rate ratio | 1 |
| Seed | 9644 |
| Sequence length | 500, 100 |
| Sequence base frequencies | Dirichlet(A=36,C=26,G=28,T=32) |
| Sequence transition rates | Dirichlet(TC=16,TA=3,TG=5,CA=5,CG=6,AG=15) |
| Controlling Duplication and Loss Rates ($5 \times 4$ conditions) | |
| Duplication rate | 4.9e-10, 2.7e-10, 1.9e-10, 5.2e-11, 0 |
| Loss rate to duplication rate ratio | 1, 0.5, 0.1, 0 |
| Controlling Duplication and ILS Rate ($3 \times 4$ conditions) | |
| Duplication rate | 4.9e-10, 1.9e-10, 0 |
| Haploid effective population size | 4.7e+8, 1.9e+8, 4.8e+7, 1e+4 |
| Controlling $n$ | |
| Number of leaves | 10, 25, 100, 250, 500 + an outgroup |
| Controlling $k$ | |
| Locus trees | 25, 100, 250, 1000, 2500, 10000 |

## C Tables

**Table S2:** Rank of methods on S100 dataset over all 120 test conditions. Ranks are obtained using mean species tree error, rounded to two significant digits to create tie for cases where error values are extremely close.

|  | 1st | 2nd | 3rd | 4th |
| --- | --- | --- | --- | --- |
| MulRF | 42 | 67 | 10 | 1 |
| DupTree | 28 | 8 | 15 | 69 |
| A-Pro | 105 | 14 | 1 | 0 |
| ASTRAL-multi | 12 | 14 | 71 | 23 |

## D Figures

**Figure S1:**
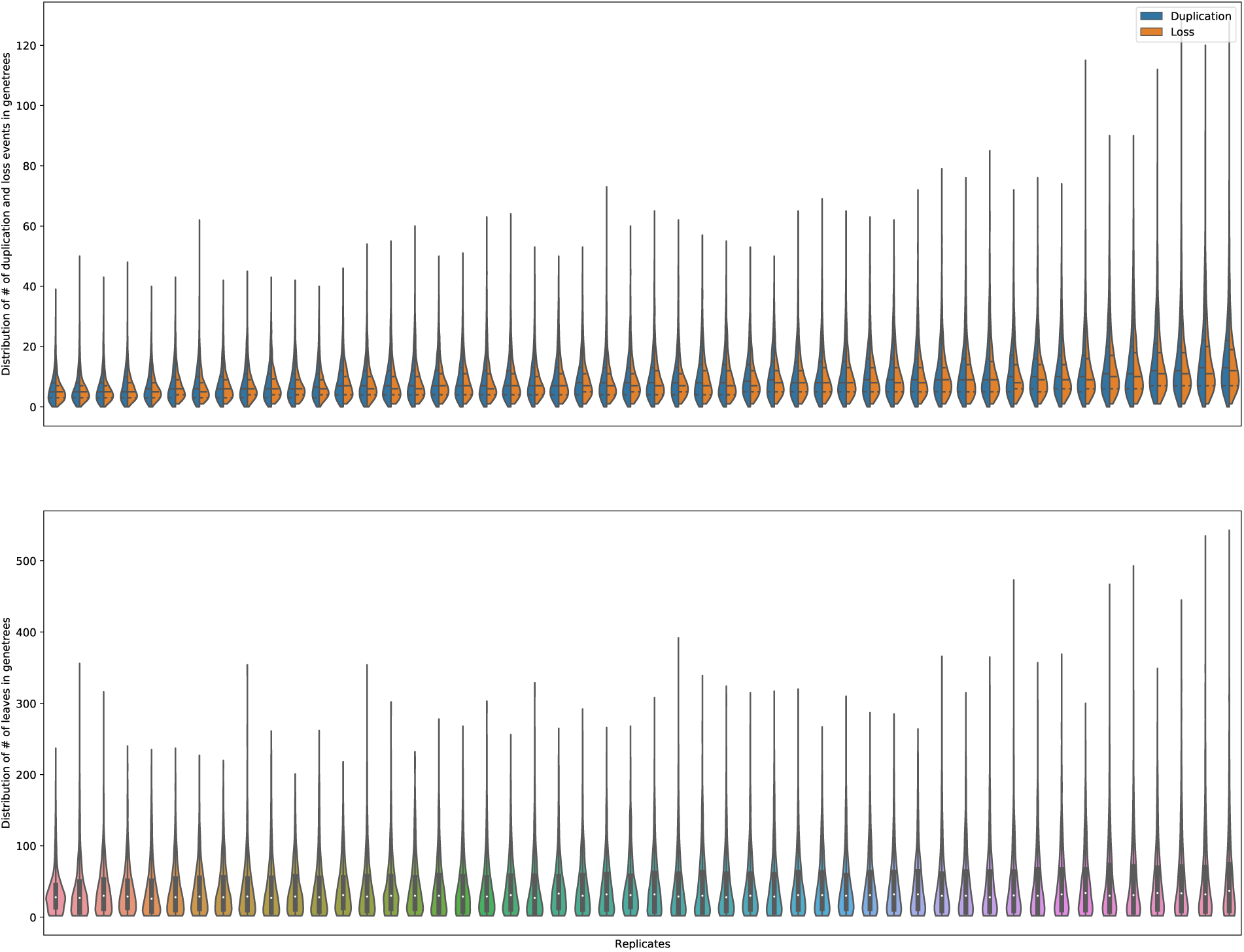
Distribution of the number of duplication events, loss events and sizes of leaf set for gene trees in the default condition by replicates. The figure on the top is sorted by the mean number of duplication events and the figure on the bottom is sorted by mean leaf set size.

**Figure S2:**
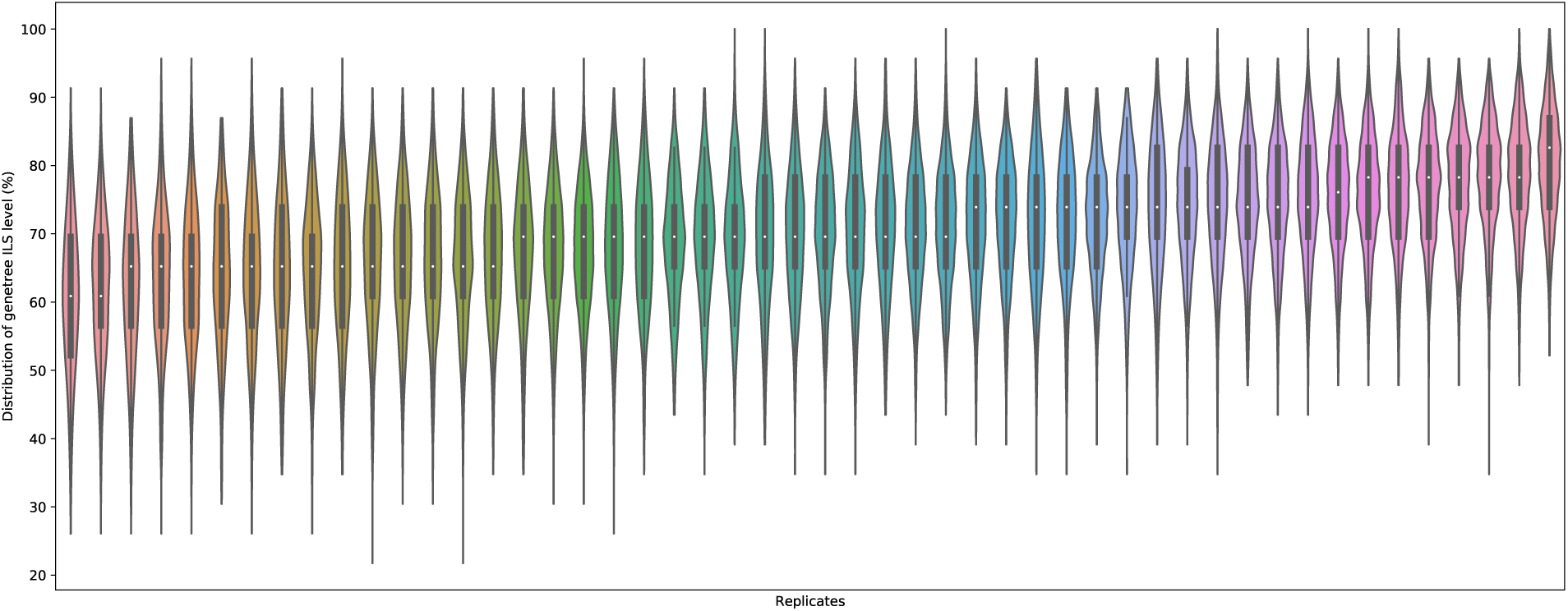
Distribution of genetree ILS, as measured by the normalized RF distance between true gene trees and the true species, in the condition with all default parameters but *λ*_+_ = *λ*_−_ = 0. Results are divided by replicates, sorted by mean ILS level.

**Figure S3:**
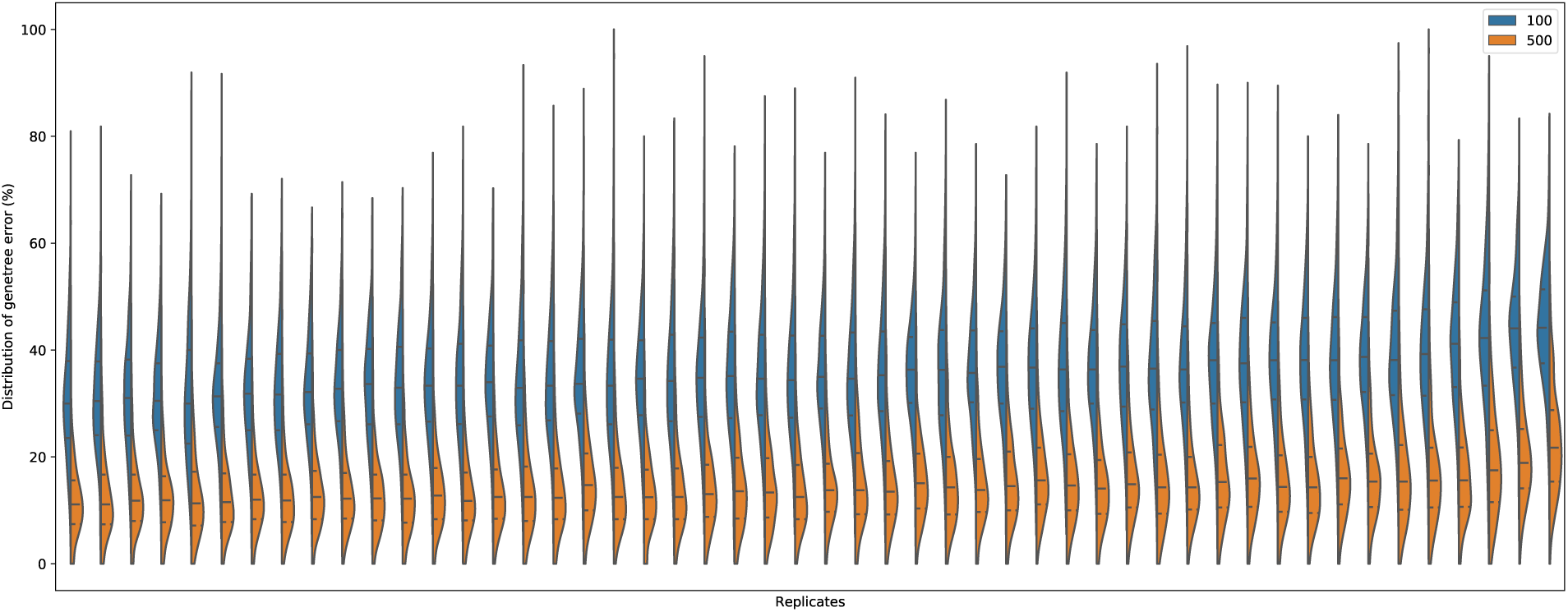
Distribution of the gene tree errors (normalized RF distance between true gene trees and the estimated gene tree) for inferred trees with at least 14 leaves in the default condition. Results are divided by sequence length (100bps or 500bps) and by replicates, sorted by mean gene tree error of the 100bps condition.

**Figure S4:**
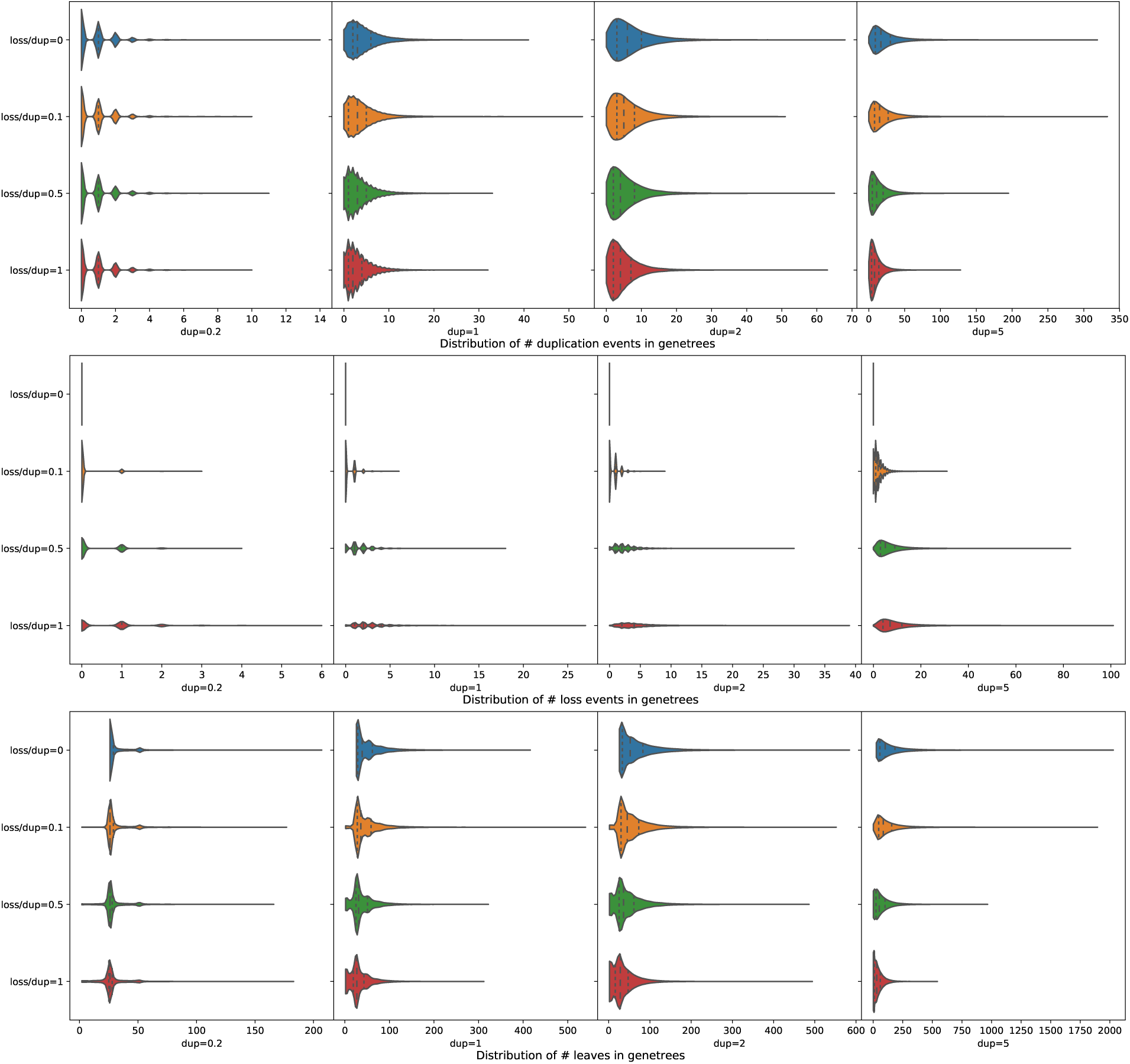
Distribution of the number of duplication events, loss events and sizes of leaf set for gene trees of each replicates sorted by duplication and loss rate.

**Figure S5:**
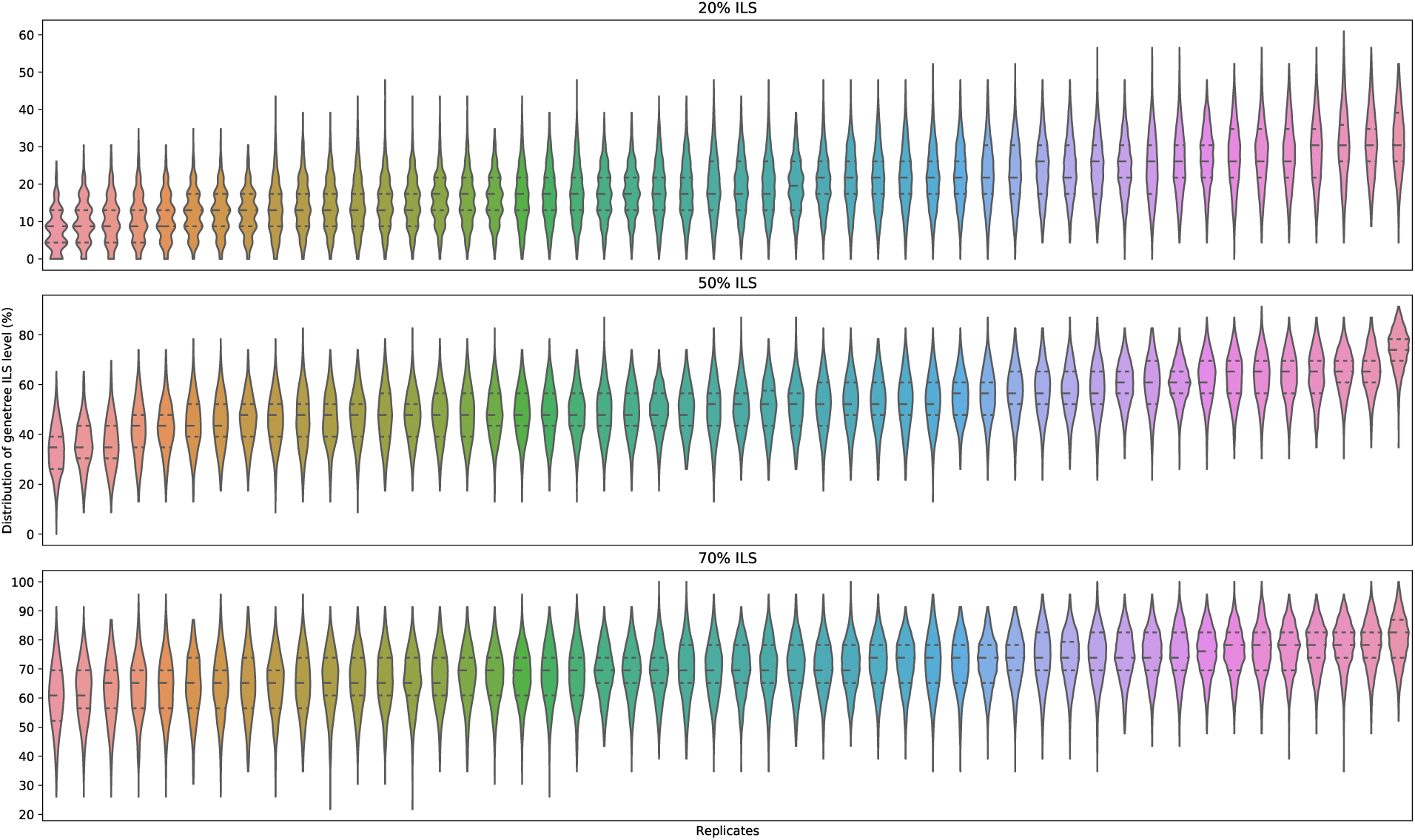
Distribution of gene tree ILS levels by replicates and expected ILS level, sorted by mean ILS level.

**Figure S6:**
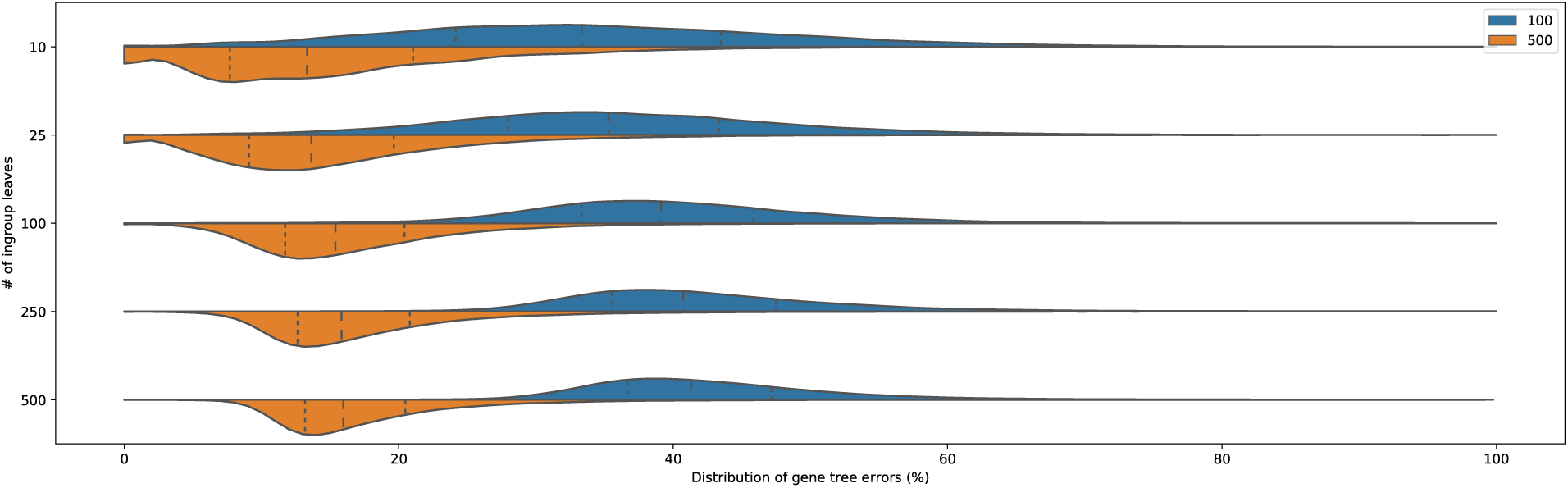
Distribution of gene tree errors by the number of ingroup species *n*.

**Figure S7:**
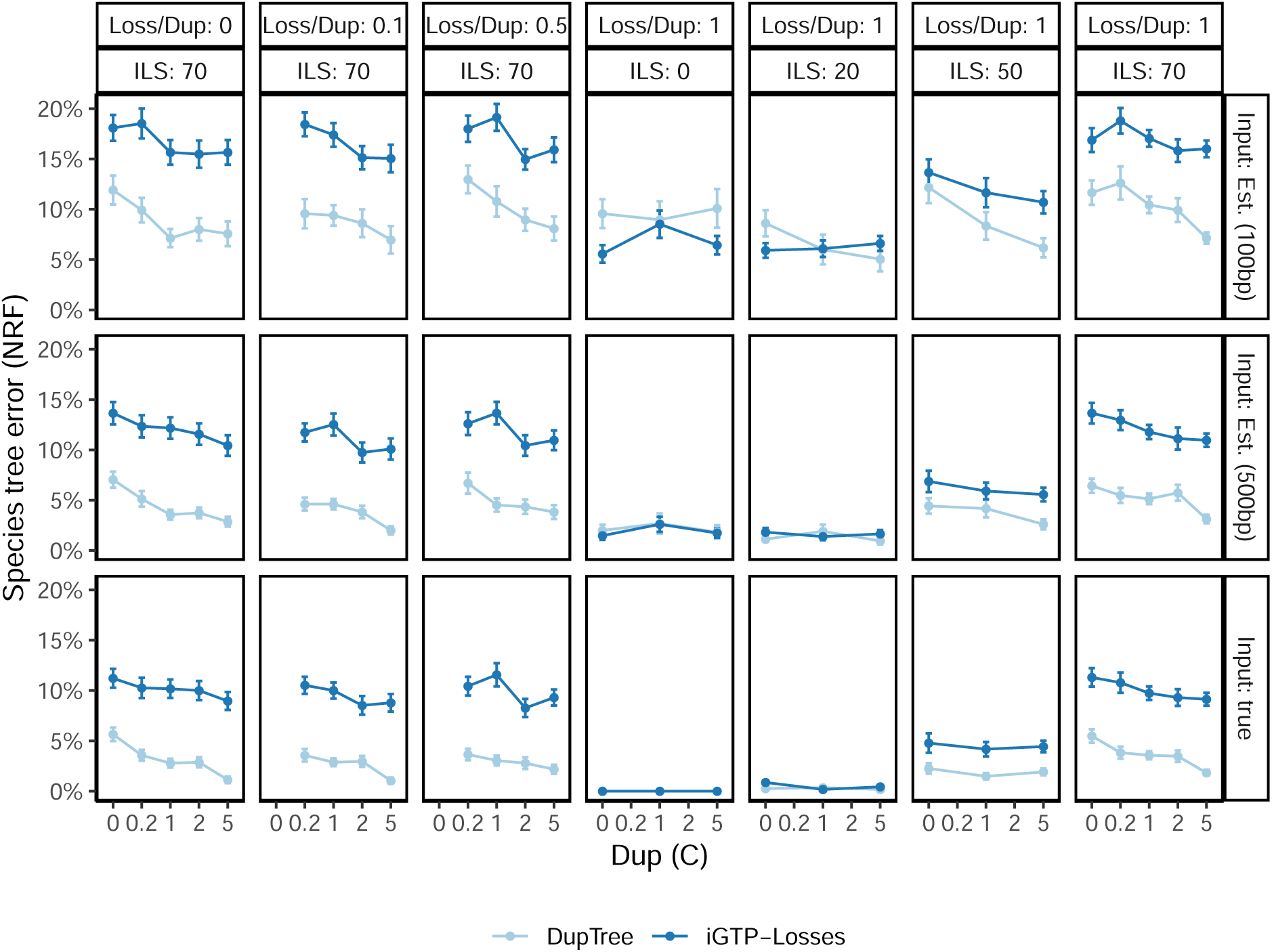
Comparison of DupTree and iGTP-DupLoss methods on all the datasets with *n* = 25 and *k* = 1000. DupTree dominates iGTP-DupLoss in most conditions.

**Figure S8:**
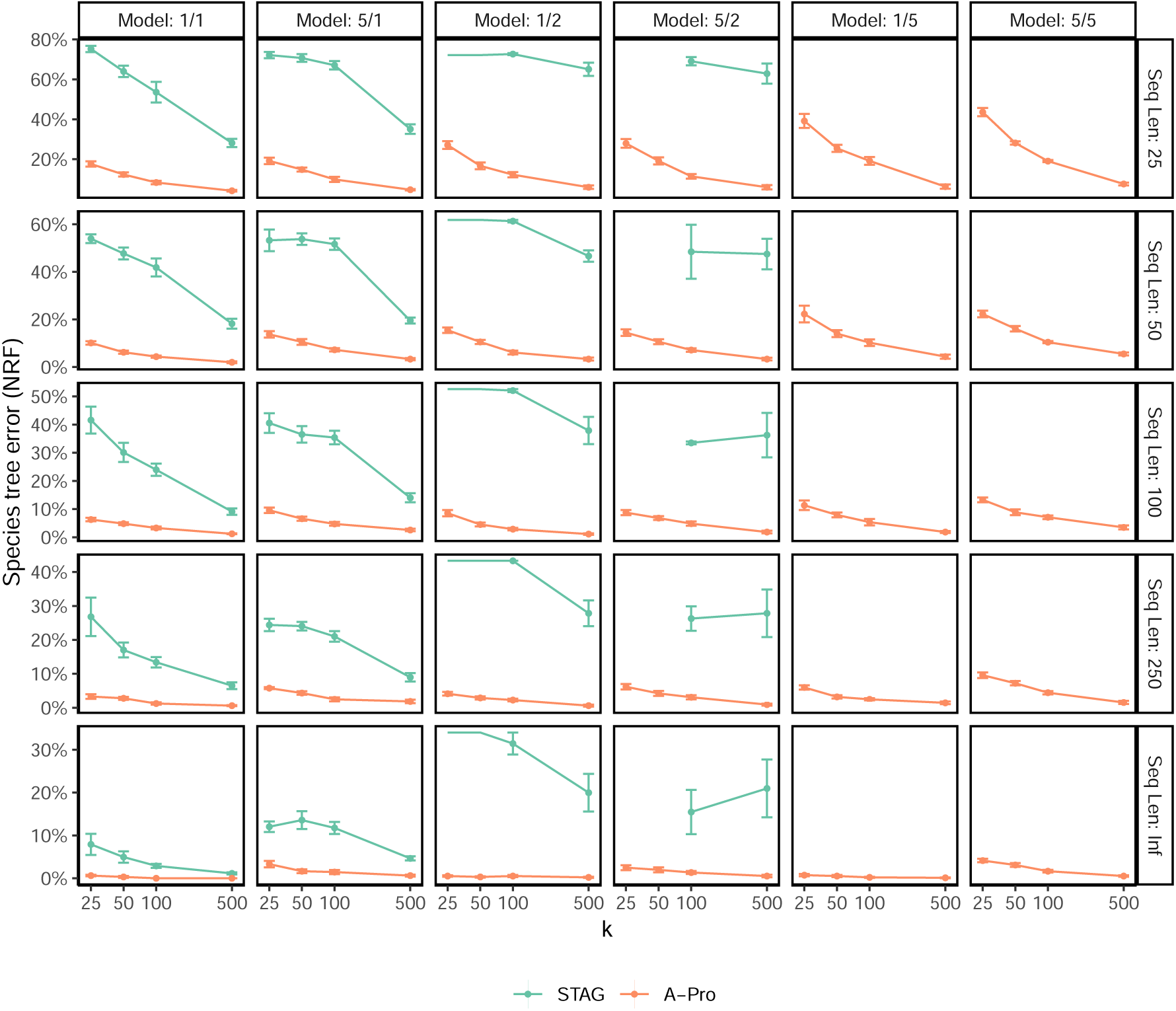
Species tree error on S100 dataset. We compare the species tree error of the STAG method to A-Pro, showing mean and standard error over 10 replicates for each model condition, with varying numbers of genes (*k*) and sequence lengths (with Inf signifying true gene trees). Model conditions are labeled as *a/b* where *a* is the level of ILS (1 or 5) and *b* is the duplication/loss rate (1, 2, or 5). Cases with missing STAG results are due to STAG failing to run on those model conditions. Note that STAG infers a species tree from the input gene trees that have at least one leaf representing each species of interest; if none of the input gene trees satisfy this requirement, then STAG fails to return a tree.

**Figure S9:**
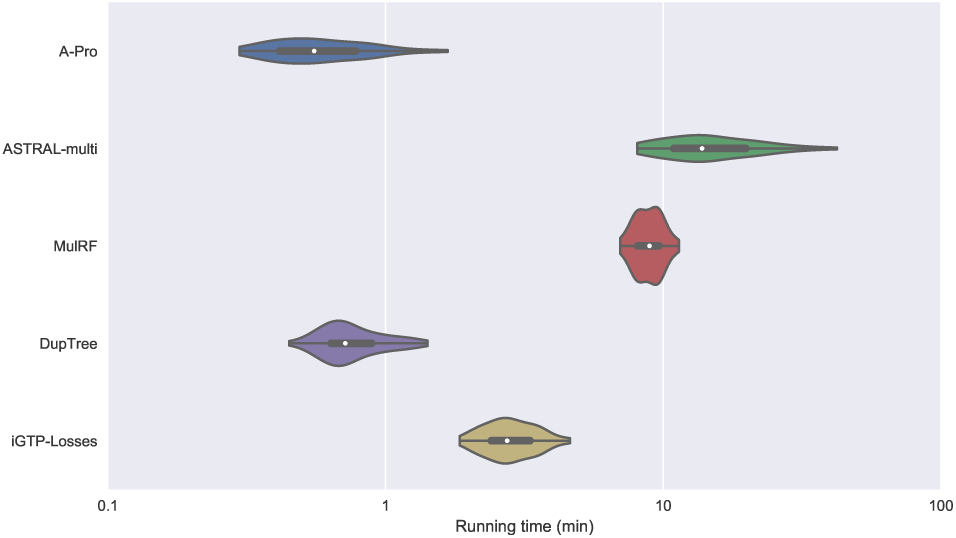
**Comparing running times**, measured on the default model condition, with estimated gene trees (100bp). All methods are run in the single-threaded mode, on the same machine with Intel(R) Xeon(R) CPU E5-2670 0 @ 2.60GHz.

**Figure S10:**
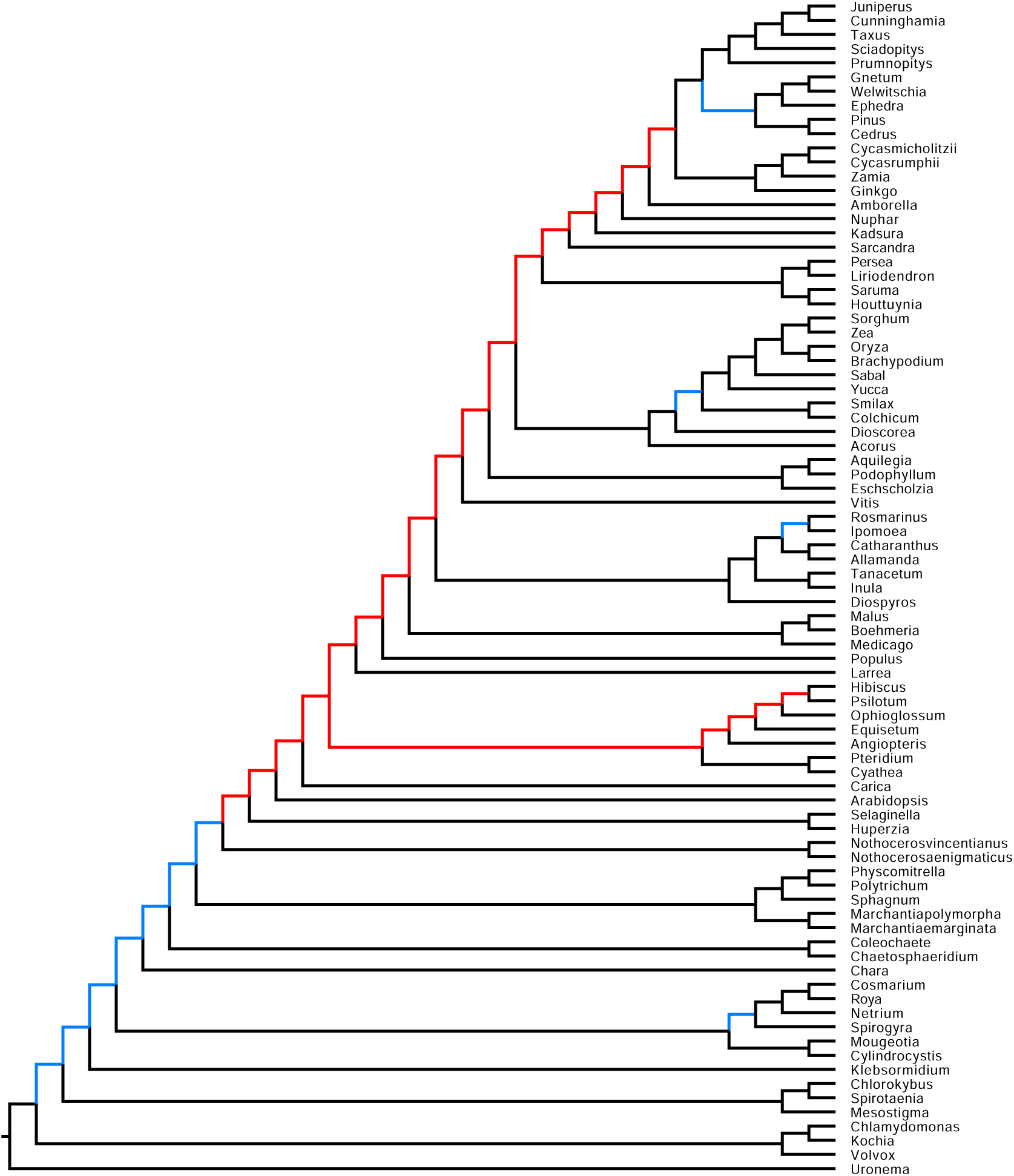
DupTree on biological plant paper. DupTree is run on 9683 multi-copy gene trees available online [56] for the plant dataset. Red: Branches that are obviously wrong, because these branches contradict basic biological categorization. Blue: Branches that contradict ASTRAL on single-copy genes that are not so obviously wrong.

